# Atherosclerosis is a smooth muscle cell-driven tumor-like disease

**DOI:** 10.1101/2023.03.06.531330

**Authors:** Huize Pan, Sebastian E. Ho, Chenyi Xue, Jian Cui, Leila S. Ross, Fang Li, Robert A. Solomon, E. Sander Connolly, Muredach P. Reilly

## Abstract

Atherosclerosis, the leading cause of cardiovascular disease, is a chronic inflammatory disease involving pathological activation of multiple cell types, such as immunocytes (e.g., macrophage, T cells), smooth muscle cells (SMCs), and endothelial cells. Multiple lines of evidence have suggested that SMC “phenotypic switching” plays a central role in atherosclerosis development and complications. Yet, SMC roles and mechanisms underlying the disease pathogenesis are poorly understood. Here, employing SMC lineage tracing mice, comprehensive molecular, cellular, histological, and computational profiling, coupled to genetic and pharmacological studies, we reveal that atherosclerosis, in terms of SMC behaviors, share extensive commonalities with tumors. SMC-derived cells in the disease show multiple characteristics of tumor cell biology, including genomic instability, replicative immortality, malignant proliferation, resistance to cell death, invasiveness, and activation of comprehensive cancer-associated gene regulatory networks. SMC-specific expression of oncogenic *Kras^G12D^* accelerates SMC phenotypic switching and exacerbates atherosclerosis. Moreover, we present a proof of concept showing that niraparib, an anti-cancer drug targeting DNA damage repair, attenuates atherosclerosis progression and induces regression of lesions in advanced disease in mouse models. Our work provides systematic evidence that atherosclerosis is a tumor-like disease, deepening the understanding of its pathogenesis and opening prospects for novel precision molecular strategies to prevent and treat atherosclerotic cardiovascular disease.

## INTRODUCTION

Atherosclerosis is the major cause of cardiovascular disease (CVD), the leading cause of death globally(*1*). It is a chronic inflammatory disease involving pathological activation of several cell types, such as immune cells (e.g., T cells, macrophages), smooth muscle cells (SMCs), and endothelial cells (ECs)(*2*). However, mechanisms underlying the disease progression and residual risk for CVD despite widespread implementation of lipid lowering therapies are unclear. Accumulating evidence suggests that smooth muscle cell (SMC) “phenotypic switching”, in which a subset of SMCs in the arterial wall proliferates, migrates, and transdifferentiates into other cell types in atherosclerotic lesions, is a central event in disease progression, lesion vulnerability, and clinical complications(*3*). Recent studies of human genetics coupled to single-cell profiling and lineage tracing have indicated that SMCs and SMC-derived cells (SDCs) contribute to the major cell phenotypes in atherosclerosis and that specific SDC sub-phenotypes can be protective or harmful in disease progression, lesion vulnerability, and clinical events(*4–7*). Even so, critical questions remain regarding behaviors and functions of SMCs in atherosclerosis and the mechanisms controlling SMC phenotypic switching.

Previous work has provided some clues on these fundamental questions. For instance, the phenomenon of SMC clonal expansion in human atherosclerosis was discovered about 50 years ago(*8*) and consolidated in studies applying multicolor lineage labeling in mouse models(*9–11*). A subset of SMCs in arterial media layer proliferates in a tumor cell-like clonal expansion manner and contributes to the cellular composition in lesions. Like tumors, DNA damage, especially oxidative DNA damage, has been widely observed in atherosclerotic lesions, especially in SMCs(*12–14*). We previously showed that several cancer-related biological processes were altered during SMC transition to “SEM” cells, a SMC-derived cell type expressing marker genes of <u>s</u>tem cells, <u>e</u>ndothelial cells, and <u>m</u>onocytes, suggesting potentially shared molecular mechanisms between atherosclerosis and tumors(*6*). Accordingly, some researchers hypothesized, at least two decades ago, that atherosclerosis represented a tumor-like arterial wall state(*15, 16*). Yet, little systematic investigation of these mechanisms has been performed to date.

Here, applying SMC lineage tracking in mouse and implementing molecular, cellular, histological, computational, genetic, and pharmaceutical studies, we reveal that atherosclerosis, in terms of SMC behaviors, share extensive commonalities with tumors. Genomic instability widely exists in both mouse and human atherosclerosis, especially in SMC lineage cells. SDCs in atherosclerosis harbor multiple tumor cell-like features and mechanistically, comprehensive cancer-associated gene regulatory networks are activated in SDCs. As a proof of concept, oncogenic mutation, *Kras^G12D^*, conditionally expressed in SMCs, promotes oxidative DNA damage, accelerates SMC phenotypic switching, and exacerbates atherosclerosis. We further demonstrate that niraparib, a clinically used anti-cancer drug targeting DNA damage repair, exhibits both preventive and therapeutic effects on atherosclerosis in mouse models. Our work represents a systematic interrogation of the pathogenesis of atherosclerosis and opens prospects for novel mechanism-based targeted strategies to prevent and treat atherosclerotic CVD.

## RESULTS

### DNA damage is associated with SMC phenotypic switching during atherosclerosis progression

Although prior studies have found that DNA damage, particularly oxidative DNA damage, is prominent in atherosclerotic lesions(*13, 14*), the link between DNA damage and SMC phenotypic switching in atherosclerosis is poorly understood. To address this, we employed SMC lineage tracing mice on low-density lipoprotein receptor knockout (*Ldlr^-/-^*) background (*ROSA26^LSL-ZsGreen1/+^; Ldlr^-/-^; Myh11-CreER^T2^*) (*6*), in which SMCs and their progenies (i.e., SDCs) were permanently labeled with ZsGreen1 in disease lesions after tamoxifen and western diet (WD) induction (**Fig. S1A**). Compared to cells in healthy arteries, 4-Hydroxy-2-Nonenal (4-HNE), a marker of lipid peroxidation and oxidative stress(*17*), was markedly increased in lesion cells, particularly in ZsGreen1^+^ SMCs and SDCs (**Fig. S1B**). Oxidative DNA damage, marked by 8-hydroxy-2’-deoxyguanosine (8-OHdG)(*18*), appeared in ZsGreen1^+^ cells at early stages of SMC phenotypic switching (e.g., at 8- and 10-week WD) (**Fig. 1, A and B**) and accumulated, especially in SDCs in intima, as atherosclerosis progressed (e.g., at 12 and 16 weeks of WD) (**Fig. 1, A-C**).

**Fig. 1.**
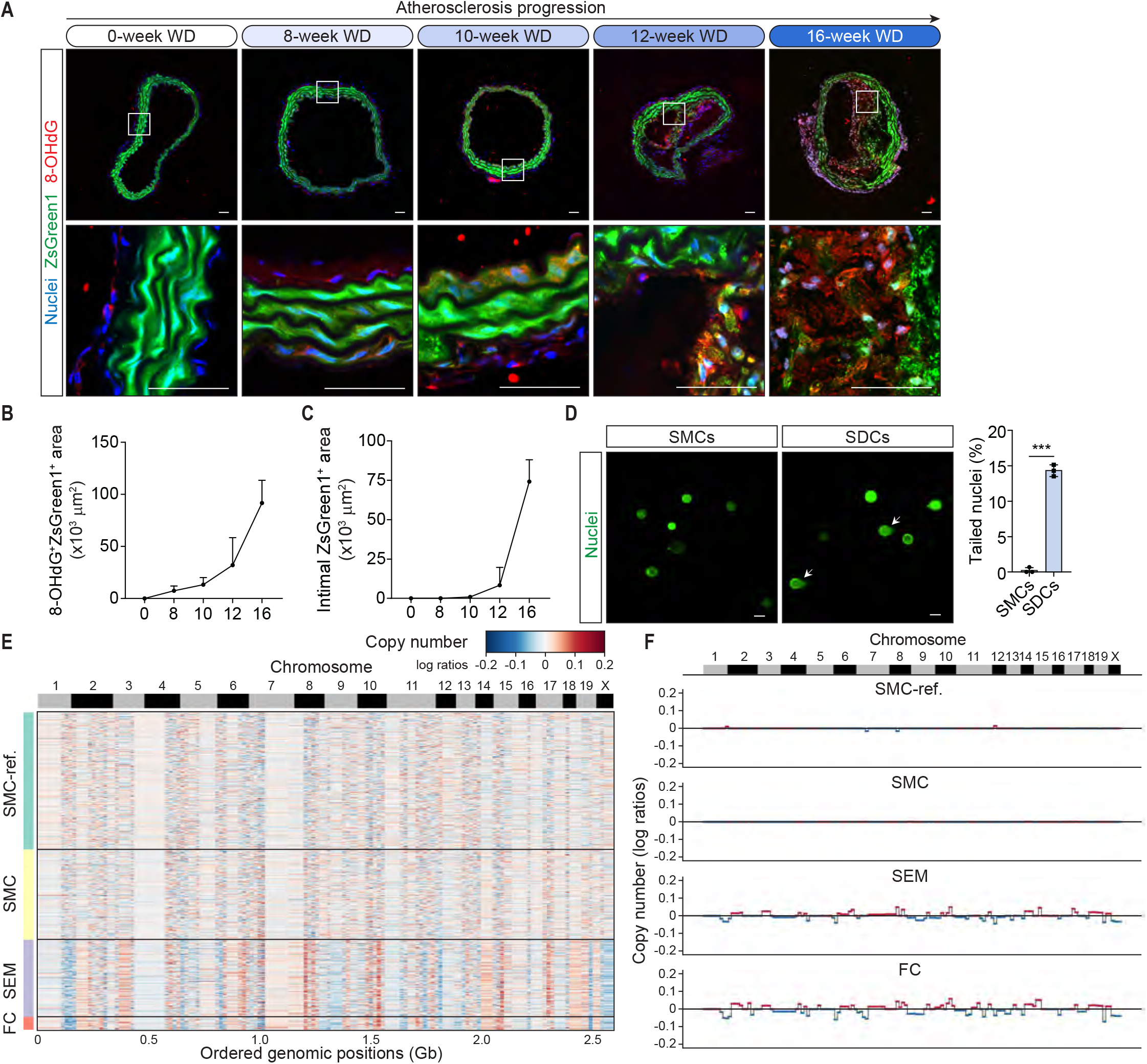
Extensive genomic instability exists in SMC lineage cells in atherosclerosis. (**A**) Immunohistochemistry (IHC) of oxidative DNA damage marker, 8-hydroxy-2’-deoxyguanosine (8-OHdG), during progression of atherosclerosis. *ROSA26^LSL-ZsGreen1/+^; Ldlr^-/-^; Myh11-CreER^T2^* mice on western diet (WD) for 0, 8, 10, 12, or 16 weeks were sacrificed for immunostaining. Representative images of brachiocephalic artery (BCA) sections from each time point are shown. (**B** and **C**) Analysis of 8-OHdG^+^ZsGreen1^+^ area in BCA sections (**B**) and ZsGreen1^+^ area in intima (**C**) suggests that oxidative DNA damage occurs at early stages of the SMC phenotypic switching process in atherosclerotic lesions. Values were indicated. N=6 mice at each time point. Scale bars, 50 μm. Significance was determined by unpaired two-tailed t test. N=3, \*\*\**P*<0.001. (**D**) Comet assay indicates tailed nuclei with single/double-strand DNA breaks. SMCs and SDCs were isolated through fluorescence-activated cell sorting (FACS) from aortas of *ROSA26^ZsGreen1/+^; Ldlr^-/-^; Myh11-CreER^T2^* mice on 0-week and 26-week WD, respectively. Statistical analysis showed the percentage of SMCs and SDCs with tailed nuclei. N=3. (**E**) Clustered heat map showing copy number profiles estimated by CopyKAT in ZsGreen1^+^ SMC lineage cell types (including SMC, SEM cell, and fibrochondrocyte (FC)) from scRNA-seq database of 16-week WD fed mice. SMCs from 0-week WD fed mice were as reference (SMC-ref.). (**F**) Line plot indicates the consensus of mouse scRNA-seq copy number profiles of each cell cluster estimated by CopyKAT in (**E**).

Sustained oxidative DNA damage can overwhelm repair pathways, leading to single- and/or double-strand DNA breaks(*19*). Indeed, estimation of tailed nuclei in the comet *assay*(*20*) revealed extensive single- /double-strand DNA breaks in SDCs pooled from advanced atherosclerosis, but very few in SMCs from healthy mouse aortas (**Fig. 1D**). Moreover, compared with isolated SMCs from healthy aortas of both young and old mice, SDCs from advanced mouse atherosclerotic lesions were markedly enriched with the doublestrand DNA break indicators(*21*), phospho-p53 (Ser15) and phospho-Histone H2A.X (Ser139) (γH2A.X) (**Fig. S1C**), suggesting that DNA damage in SDCs is an atherosclerosis-related rather than simply an aging event. Consistent with previous reports(*12, 13*), cells with DNA damage, as indicated by γH2A.X staining, were also observed in human atherosclerotic lesions (**Fig. S1D**). Taken together, these data support an association between DNA damage, especially oxidative DNA damage, and SMC phenotypic switching during atherosclerosis progression.

### Extensive genomic instability exists in SMC lineage cells of both mouse and human atherosclerosis

Accumulation of unrepaired DNA damage can induce genomic instability(*22*). To probe genome integrity in lesion cells, especially in SDCs, during atherosclerosis progression, we applied computational approaches, CopyKAT(*23*) and inferCNV(*24*), to both mouse and human atherosclerotic scRNA-seq data(*6*). These tools identify large-scale chromosomal copy number variations (CNVs) based on the genome-wide expression intensity in scRNA-seq data. Almost all cell types in mouse lesions, especially SDCs (e.g., SEM cells and fibrochondrocytes (FCs))(*6*), showed large-scale genomic CNVs (**Fig. 1E; Fig. S2, A and B**). In SEM cells and FCs, large-scale reductions in chromosomal copy numbers were found in chromosome 1, 5, 6, 9, 12, 19, and X, while increased copy numbers were observed in chromosome 2, 3, 6, 8, 10, 15, and 17 (**Fig. 1, E and F**). FCs exhibited a greater extent of CNVs than SMCs and SEM cells (**Fig. S2, C and D**), suggesting an accumulation of genomic instability along the trajectory of SMC phenotype switching. Although there was higher heterogeneity in human versus mouse atherosclerotic lesions, both CopyKAT and inferCNV analyses of human scRNA-seq data from atherosclerotic plaques of multiple patients undergoing carotid endarterectomy (CEA) revealed widespread genomic instability in lesion cells, including SMCs and putative SDCs (e.g., intermediate cell state (ICS) and FCs) (**Fig. S3, A-F**). These results indicate that extensive genomic instability exists in atherosclerosis, especially in SDCs.

### SMC-derived cells in atherosclerosis harbor multiple tumor cell-like characteristics

Depending on the cell and tissue environment, genomic instability can result in one or different combinations of senescence, apoptosis, and malignant proliferation, ultimately accelerating aging, degenerative disease, and/or cancer(*25*). The activity of senescence-associated beta-galactosidase (SA-β-Gal), a biomarker of cellular senescence(*26*), was undetectable in young mouse arteries (**Fig. S4A**, left panel), but was markedly elevated in the medial SMC layer of aged non-atherosclerotic arteries (**Fig. S4A**, middle panel). Intriguingly, most cells in the intima of disease lesion, enriched with SDCs and other cells, were negative for SA-β-Gal staining, though some fibrous cap and shoulder regions of atherosclerotic arteries were stained with SA-β-Gal (**Fig. S4A**, right panel). Furthermore, in cultured aortic SMCs from healthy young mice and SDCs from mice with advanced atherosclerotic lesions (**Fig. S4B**), about 30% of primary SMCs at passage 5 (P5) were positive for SA-β-Gal, while almost no SDCs at this passage or much later (e.g., up to P45) was SA-β-Gal^+^ (**Fig. 2A**). Moreover, expression of two cell senescence marker proteins, p21 and p16(*27*), were decreased in *ex vivo* SDCs versus SMCs (**Fig. 2B**). These results suggest that SDCs in atherosclerosis have largely escaped from replicative senescence.

**Fig. 2.**
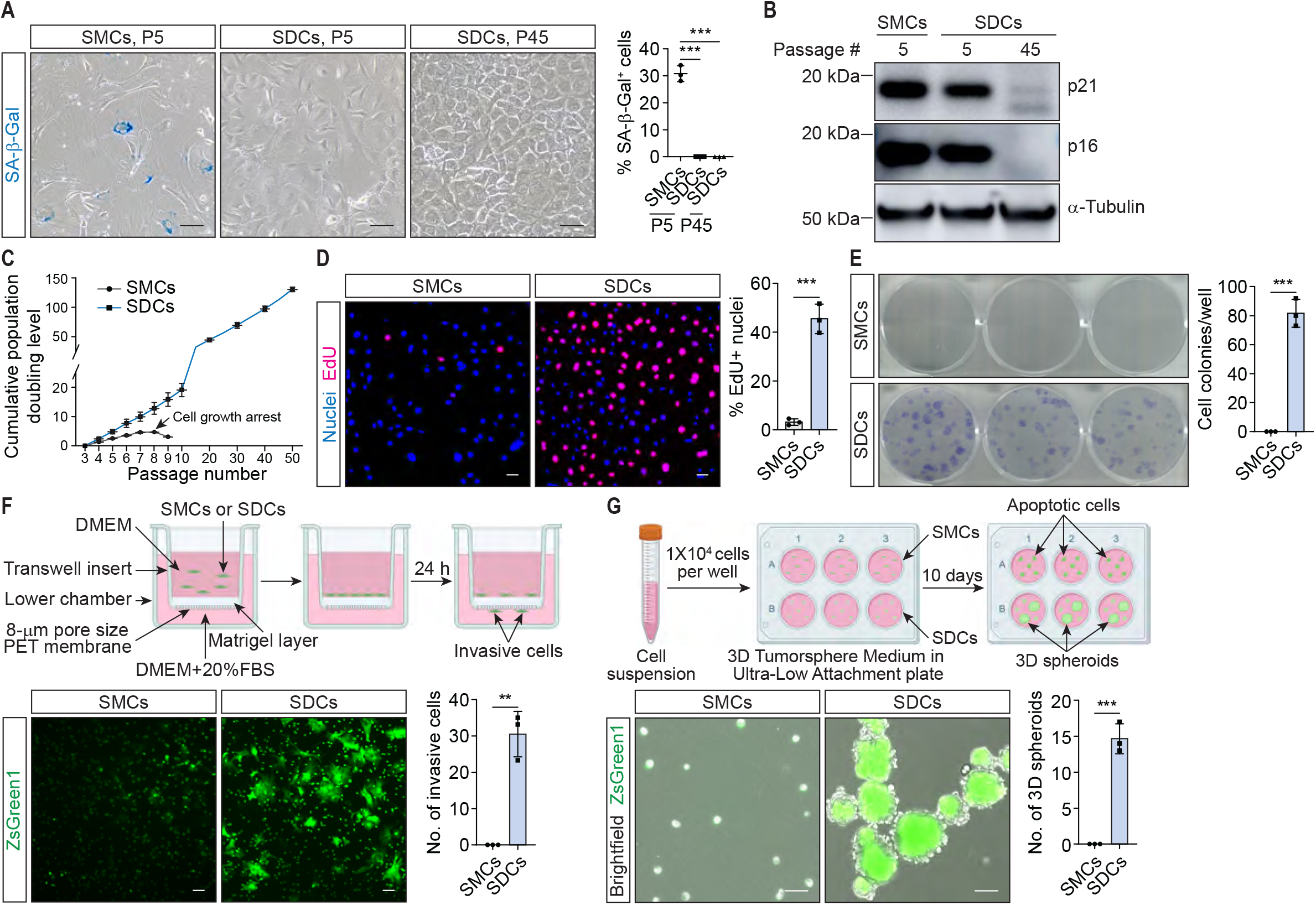
SMC-derived cells formed in atherosclerosis show multiple tumor cell-like characteristics. (**A**) Staining of senescence-associated beta-galactosidase (SA-β-Gal), a biomarker of cellular senescence, in *ex vivo* SMCs (at passage 5 (P5)) and SDCs (at P5 and P45). Proportion of SA-β-Gal^+^ SMCs or SDCs at each passage was analyzed. (**B**) Immunoblotting of cellular senescence markers, p21 and p16, indicates that senescence marker proteins are reduced in SDCs at early passage (P5) and late passage (P45) compared to SMCs at P5. (**C**) Calculation of cell population doubling levels of SMCs and SDCs starting from passages 3 indicates that SDCs harbor replicative immortality while SMCs encounter growth arrest within 10 passages. (**D**) SMCs and SDCs were incubated with 5-ethynyl-2’-deoxyuridine (EdU) for 2 hours. Click-iT EdU assay indicated that SDCs had much higher proliferative rate (proportion of EdU^+^ nuclei) than SMCs. (**E**) *Ex vivo* SDCs form colonies at low seeding density (200/well, 6-well plate). Cell colonies per well were counted. (**F**) Schematic and representative images of cell invasion assay with SMCs and SDCs. Number of cells invading through Matrigel layer were shown. (**G**) Schematic and representative images of 3D spheroid formation assay with SMCs and SDCs. SMCs and SDCs were seeded into ultra-low attachment plate and cultured with 3D Tumorsphere Medium for 10 days. 3D spheroids were counted. Scale bars, 50 μm. Significance was determined by unpaired two-tailed t test. N=3, \*\**P*<0.01, \*\*\**P*<0.001.

*Ex vivo* lesion SDCs also exhibited replicative immortality, with all cultured SDC cell lines showing no sign of senescence or slower proliferation even at advanced passages (e.g., >50 passages) (**Fig. 2C**). In contrast, isolated SMCs underwent growth arrest within 10 passages (**Fig. 2C**). This suggests that SDCs in atherosclerosis, unlike normal primary cells in culture, are not subjected to the Hayflick Limit(*28*) but have acquired malignant proliferative capacity like tumor cells. Indeed, SDCs had much higher proliferation rate than SMCs, as indicated by markedly increased integration of 5-ethynyl-2’-deoxyuridine (EdU)(*29*) into newly synthesized DNA in SDCs (**Fig. 2D; Fig. S4C**). Compared with SMCs, individual SDCs were also able to grow into large colonies at low cell seeding density (**Fig. 2E**) and showed much lower baseline and TNFα-induced cell death (**Fig. S4D**). These data suggest that SDCs resist cell death and have significantly enhanced cell survival. Like most tumor cells(*30*), SDCs also acquired invasiveness, indicated by the capacity of invading through extracellular matrix (**Fig. 2F**). In a serum-free suspension culture system(*31*), some *ex vivo* SDCs formed three-dimensional (3D) spheroids, exhibiting a characteristic of cancer stem cells (**Fig. 2G**). In fact, SDCs from atherosclerotic plaques had increased expression of cancer stem cell marker genes(*32*), such as CD24 and CD44 (**Fig. S4, E-G**). Overall, these data suggest that DNA damage-induced genomic instability in atherosclerotic lesion results in SDCs with multiple tumor cell-like characteristics.

### Atherosclerosis shares comprehensive gene regulatory networks with cancer

To define molecular regulatory networks underlying tumor cell-like phenotypic change of SMCs in atherosclerosis, we employed Pathway RespOnsive GENes analysis (PROGENy)(*33*) to infer pathway activity based on mouse atherosclerotic scRNA-seq database and validated the predicted results for each signaling pathway. The activities of 12 major cancer-associated signaling pathways(*33*), including EGFR, Hypoxia, JAK-STAT, MAPK, NFκB, p53, PI3K, TNFα, Trail, VEGF, and WNT, except TGFβ, were predicted to be activated in SDCs (i.e., SEM cells and FCs) compared with SMCs (**Fig. 3A; Fig. S5A**).

**Fig. 3.**
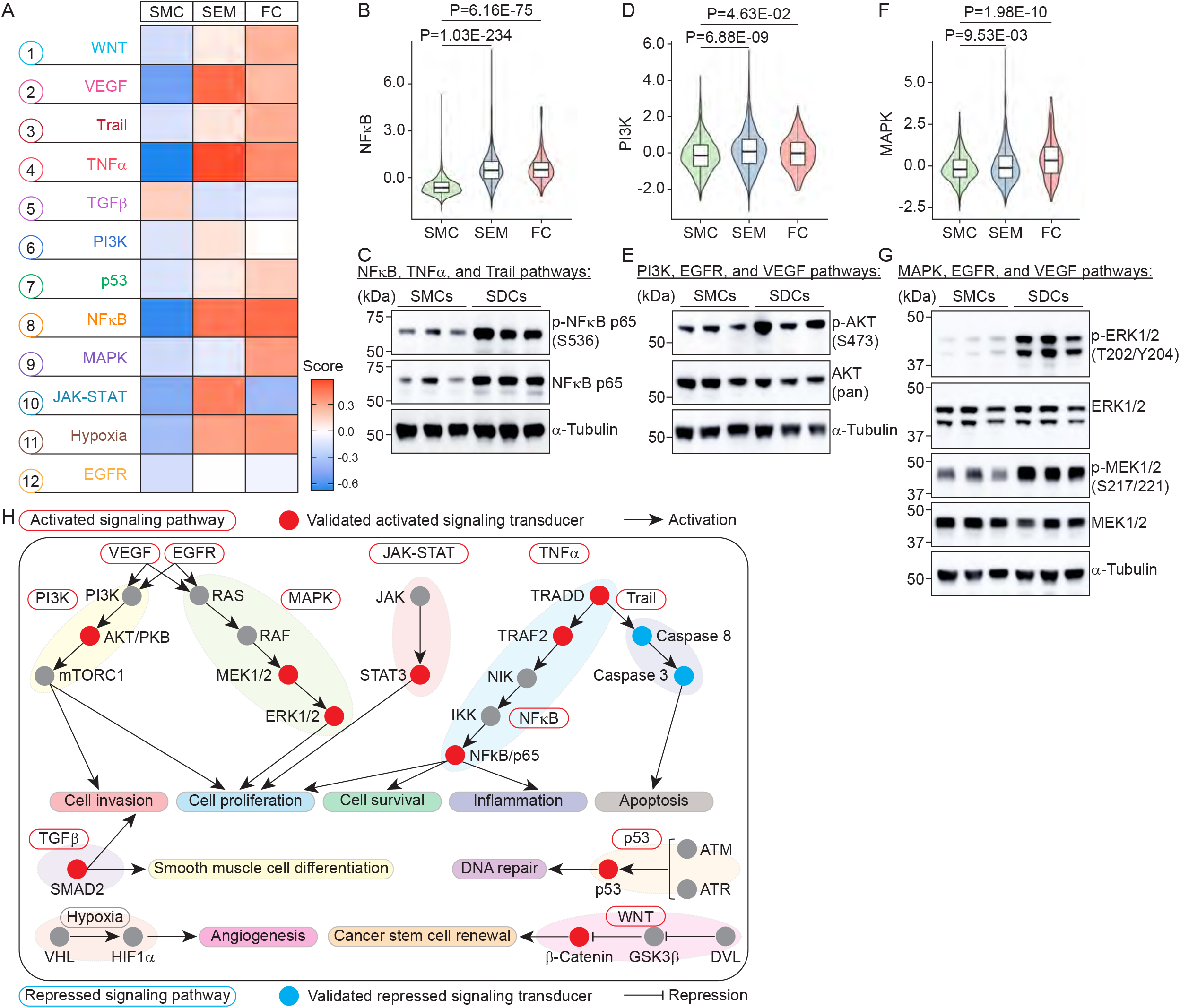
Analysis of cancer-associated signaling pathways activated in SDCs versus SMCs in atherosclerosis. (**A**) Heat map showing median scores of 12 cancer-associated signaling pathways in SMCs, SEM cells, and FCs estimated by Pathway RespOnsive GENes (PROGENy). (**B**) Violin plot shows PROGENy score for NFκB pathway. (**C**) Immunoblotting results indicate that phospho-NFκB p65 (S536) was increased in SDCs versus SMCs. (**D**) Violin plot shows PROGENy score for PI3K pathway. (**E**) Immunoblotting results indicate that phospho-AKT (S473) was increased in SDCs versus SMCs. (**F**) Violin plot shows PROGENy score for MAPK pathway. (**G**) Immunoblotting results indicate that phospho-ERK (T202/Y204) and phospho-MEK1/2 (S217/221) were increased in SDCs versus SMCs. (**H**) Summary of 12 cancer-prone signaling pathways that were activated in SDCs. Key transducers of each signaling pathway that were validated via immunoblotting were marked by red (activated) or blue (repressed) dots. Cancer-related functions of each signaling pathway were indicated. *P* values are shown.

Enhanced p53 pathway(*34*) activity (**Fig. S1C, S5B**) suggests that DNA damage response is induced in SDCs during atherosclerosis progression, consistent with elevated DNA damage (**Fig. 1A**) and genomic instability (**Fig. 1E**) in SDCs. Activation of multiple signaling pathways in SDCs, including inflammatory response-related (e.g., TNFα(*35*) (**Fig. S5, C and D**) and NFκB(*36*) (**Fig. 3, B and C**)), PI3K(*37*) (**Fig. 3, D and E**), MAPK(*38*) (**Fig. 3, F and G**), EGFR(*39*) (**Fig. 3, E and G; Fig. S5E**), VEGF(*40*) (**Fig. 3, E and G; Fig. S5F**), and JAK-STAT(*41*) signaling (**Fig. S5, G and H**), may account for the marked enhancement of SDC proliferation (**Fig. 2, C and D**). Activation of NFκB(*36*) (**Fig. 3, B and C**) and Trail(*42*) (**Fig. 3C; Fig. S5I**) in SDCs may promote cell survival (**Fig. 2E**) and underlie SDC resistance to induced cell death (**Fig. S4D**). Elevated PI3K(*37*) activity (**Fig. 3, D and E**) could also be responsible for the invasiveness of SDCs (**Fig. 2F**). Activation of other cancer-related signaling pathways (e.g., WNT(*43*) (**Fig. S5, J and K**)) may contribute to the cancer stem cell-like renewal of SDCs (**Fig. 2G**). As an exception, TGFβ signaling, a key promotor of SMC differentiation(*44*), was predicted to be suppressed during SMC phenotypic switching (**Fig. S5L**) but was found to be induced in *ex vivo* SDCs (**Fig. S5M**). These data demonstrate that during progression of atherosclerosis, the phenotypic alteration of SMCs is associated with a comprehensive transformation to gene regulatory networks resembling that in tumor cells (**Fig. 3H**).

### Kras^G12D^, an oncogenic mutation, accelerates SMC phenotypic switching and atherosclerosis progression

To further interrogate whether tumor-like pathogenesis can induce SMC phenotype switching and drive atherosclerosis progression, we introduced oncogenic *Kras^G12D^* to the SMC lineage tracing mice (**Fig. S6A**). The KRAS^G12D^ mutant was chosen because *KRAS* is one of the most commonly mutated genes associated with progression of multiple types of cancer(*45*), the oncogenic KRAS^G12D^ was reported to drive tumorigenesis through oxidative DNA damage(*46*) and genomic instability, and in particular, constitutively activated KRAS (e.g., KRAS^G12D^) triggers at least 3 cancer-associated signaling pathways, including NFʺB, PI3K, and *MAPK*(*47–49*), which were hyperactivated in SDCs (**Fig. 3, B-G**). SMC-specific expression of *Kras^G12D^* had no significant influence on body weight (**Fig. S6B**) or serum cholesterol level (**Fig. S6C**), but *Kras^LSL-G12D/+^; ROSA26^LSL-ZsGreen1/+^; Ldlr^-/-^; Myh11-CreER^T2^* (hereafter SMC-*Kras^G12D/+^*) mice had increased atherosclerotic lesion area compared with SMC-*Kras^+/+^* control mice (**Fig. S6, D and E**), suggesting that *Kras^G12D^* specifically expressed in SMCs can exacerbate atherosclerosis.

In the time course study, in which mice were sacrificed at 0, 8, 10, and 12-week WD (**Fig. 4A**), SMC-specific *Kras^G12D^* expression markedly increased SMC lineage cells with oxidative DNA damage (8-OHdG^+^ZsGreen1^+^) (**Fig. 4, A and B**) and total SMC lineage cells (ZsGreen1^+^) (**Fig. 4C**) during progression of the disease, starting from 8-week WD feeding. Compared with the control counterparts, SMC-*Kras^G12D/+^* mice showed larger atherosclerotic lesion areas at each time point of WD (**Fig. 4D**). Like tumor development, SMC phenotypic switching exhibits phased dynamics, which is consistent with previous reports(*10*), and can accordingly be divided into four stages: i) original contractile SMC (e.g., vascular SMCs of mice at 0-week WD), ii) early remodeled SMC/SDC (e.g., at 8-week and 10-week WD), iii) fibrous cap SMC/SDC (e.g., at 12-week WD), and iv) neointimal SDC (e.g., at 16-week WD) (**Fig. 1A**). As indicated in **Fig. 4E**, at each time point of WD, a higher proportion of *SMC-Kras^G12D/+^* mice compared with the control mice developed more advanced SMC phenotypic switching, from contractile SMC to early remodeled lesion SMC/SDC to fibrous cap SMC/SDC to neointimal SDC. Altogether, these results provide a proof of concept that SMC oncogenic mutations can accelerate atherosclerosis progression by inducing SMC phenotypic switching.

**Fig. 4.**
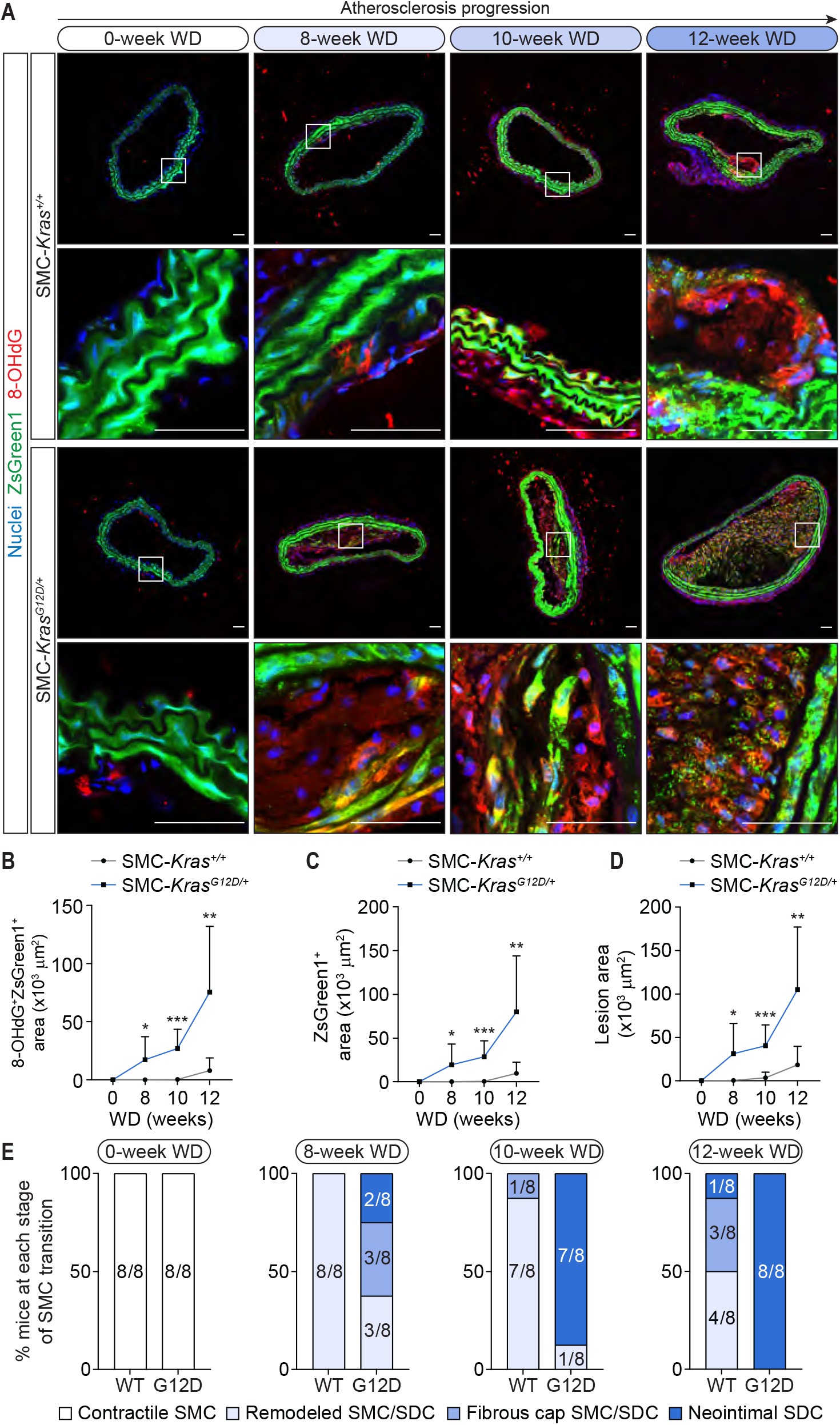
SMC-specific expression of *Kras^G12D^* accelerates SMC phenotypic switching during atherosclerosis progression. (**A**) *Kras^+/+^; ROSA26^LSL-ZsGreen1/+^; Ldlr^-/-^; Myh11-CreER^T2^* (SMC-*Kras^+/+^*) and *Kras^LSL-G12D/+^; ROSA26^LSL-ZsGreen1/+^ Ldlr^-/-^; Myh11-CreER^T2^* (SMC-*Kras^G12D/+^*) mice were sacrificed for IHC staining after 0, 8, 10, or 12 weeks of WD. Representative images of mouse BCA sections stained with oxidative DNA damage marker, 8-OHdG, at each time point are shown. (**B**-**D**) Statistical analysis of 8-OHdG^+^ZsGreen1^+^ area (**B**) and ZsGreen1^+^ area (**C**) in neointima and total atherosclerotic lesion area (**D**) in BCA sections. (**E**) SMC phenotypic switching was divided into four stages: (i) contractile SMC, (ii) early remodeled SMC/SDC, (iii) fibrous cap SMC/SDC, and (iv) neointimal SDC. Proportion and number of SMC-*Kras^+/+^* and SMC-*Kras^G12D/+^* mice at each stage of SMC phenotypic switching at each time point are indicated. N=8 mice/group at each time point. Scale bars, 50 μm. Significance was determined by unpaired two-tailed t test. \**P*<0.05, \*\**P*<0.01, \*\*\**P*<0.001.

### Niraparib, a clinical anti-cancer drug targeting DNA damage repair, shows beneficial effects on atherosclerosis

We then explored if certain anti-cancer therapies, such as strategies targeting DNA damage observed in atherosclerotic SDCs, could be beneficial to the disease. Niraparib, a poly(adenosine diphosphate–ribose) polymerase (PARP) inhibitor suppressing DNA damage repair, is an FDA-approved chemotherapy for various cancer types(*50*). Administration of niraparib to atheroprone mice during atherosclerosis progression (**Fig. 5A**) had few effects on mouse body weight (**Fig. S7A**) or circulating cholesterol levels (**Fig. S7B**). Niraparib significantly attenuated SMC phenotypic switching and reduced SDCs with oxidative DNA damage (8-OHdG^+^ZsGreen1^+^) (**Fig. S7, C and D**) as well as total SMC lineage cells (ZsGreen1^+^) (**Fig. S7, C and E**). Further, niraparib markedly decreased atherosclerotic lesion area (**Fig. 5, A and B**) and increased features of atherosclerotic plaque stability by reducing necrotic core area (**Fig. 5C**) and increasing the ratio of fibrous cap/lesion area (**Fig. 5D**). In mice with established atherosclerosis (mice on 16-week WD), niraparib administration during an additional 8 weeks of WD reduced atherosclerotic plaque burden in the whole aorta (**Fig. 5, E and F**) and lesion area in aortic sinus (**Fig. 5, G and H**) as well as improved lesion stability by decreasing necrotic core area (**Fig. 5I**) and increasing ratio of fibrous cap/lesion area (**Fig. 5J**). These findings suggest that niraparib, an anti-cancer chemotherapy targeting DNA damage, also has both preventive and therapeutic benefits on atherosclerosis and features of plaque stability in mouse models.

**Fig. 5.**
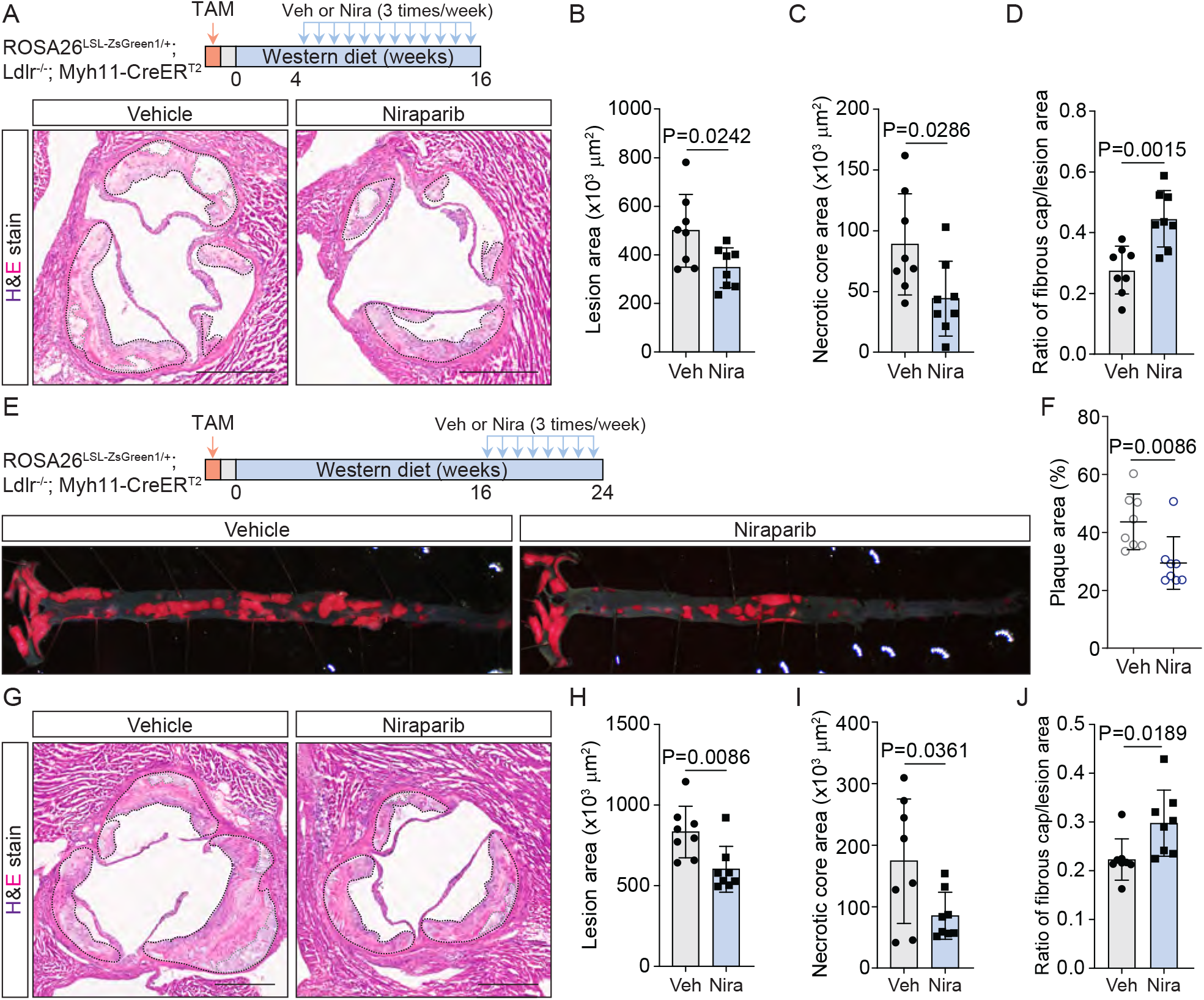
Niraparib has both preventive and therapeutic effects on atherosclerosis in mouse models. (**A**) Schematic of administration of niraparib to *ROSA26^LSL-ZsGreen1/+^; Ldlr^-/-^; Myh11-CreER^T2^* mice during initiation and progression of atherosclerosis. The mice were induced with tamoxifen (TAM) for one week and then fed one-week chow diet, followed by WD. After 4 weeks of WD, mice were treated with niraparib (Nira, 10 mg/kg mice, 3 times/week) or vehicle (Veh, corn oil, 3 times/week) and sacrificed after 16-week WD. Aortic sinus sections were subjected to hematoxylin and eosin (H&E) stain. Representative images from Veh and Nira-treated mice were shown. (**B**-**D**) Statistical analysis of lesion area (**B**), necrotic core area (**C**), and ratio of fibrous cap/lesion area (**D**) in H&E-stained sections from Veh and Nira-treated mice in (**A**). (**E**) Schematic of niraparib treatment to *ROSA26^LSL-ZsGreen1/+^; Ldlr^-/-^; Myh11-CreER^T2^* mice with established atherosclerosis. The mice were induced with TAM and fed chow diet followed by WD as the same in (**A**). After 16 weeks of WD, mice were treated with Nira (10 mg/kg mice, 3 times/week) or Veh (corn oil, 3 times/week) and sacrificed at total of 24-week WD. *En face* aortas were stained with Oil Red O and captured under microscope. Representative images from mice of two groups were shown. (**F**) Proportions of plaque area in aortas from Veh and Nira-treated mice in (**E**). (**G**) Representative images of H&E-stained aortic sinus sections from Veh and Nira-treated mice with established atherosclerosis were shown. (**H**-**J**) Statistical analysis of lesion area (**H**), necrotic core area (**I**), and ratio of fibrous cap/lesion area (**J**) in H&E-stained sections from Veh and Nira-treated mice in (**G**). Scale bars, 500 μm. Significance was determined by unpaired two-tailed t test. N=8 mice/group. *P* values are indicated.

## DISCUSSION

Based on our novel findings centered on the SMC transformation in atherosclerosis, we propose “athero-oncology” as an innovative integrative concept for fundamental research and translational focus of atherosclerotic CVD. SMC phenotypic switching in atherosclerosis shares extensive commonalities with tumor biology in terms of genomic instability, tumor cell-like characteristics of SDCs, activation of cancer-associated signaling pathways, oncogene-driven SMC transformation and disease modulation, and response to anti-cancer treatments that target DNA damage (**Fig. S8**). Our study further endorses the prominent roles of SMCs and SDCs in atherosclerosis and provides a novel mechanistic lens for the development of precision medicine for atherosclerotic CVD.

Our current work advances previous findings of DNA damage in atherosclerotic lesions(*13, 14*) by uncovering extensive genomic instability in both mouse and human atherosclerosis, particularly in lesion SDCs. Instead of promoting cellular senescence, it appears that genomic instability triggers escape of senescence, and subsequently replicative immortality and tumor cell-like transformation of SMCs in atherosclerosis. In fact, a paucity of senescent cells in atherosclerotic lesions was observed in advanced human atherosclerosis(*51*). Clearance of senescent cells in intima and at the fibrous cap could attenuate atherosclerosis(*52*) and enhance lesion stability by promoting SMCs towards a promigratory phenotype(*53*). Our proof of principle genetic and pharmacological studies indicates that combinations of SMC-targeted chemotherapies and senotherapies may provide new opportunities to improve lesion stability particularly in the context of residual CVD risk in patients already on lipid lowering therapies.

Tumor cell-like genomic instability may be one of driving factors for SMC transformation in atherosclerosis. Indeed, Bennett group showed that enhancing repair of double strand DNA breaks or oxidative DNA damage by SMC-specific overexpression of nijmegen breakage syndrome 1 (NBS1)(*13*) or 8-oxoguanine DNA glycosylase (OGG1)(*14*), respectively attenuated and stabilized atherosclerotic lesions in mouse models. It remains unclear, however, whether specific genomic mutations trigger and drive SMC tumor-like transformation in atherosclerosis as that occurs in tumorigenesis. As a proof of concept, we demonstrate that SMC conditional expression of oncogenic *Kras^G12D^* promotes oxidative DNA damage, accelerates SMC phenotypic switching, and exacerbates atherosclerosis. Previously, it was shown that deficiency of the tumor suppressor, *Trp53*, resulted in larger atherosclerotic lesion and increased proliferation and reduced apoptosis in several cell types (e.g., SMCs, macrophages) in atherosclerosis(*54*). Indeed, clonal hematopoiesis of indeterminate potential, the age-related expansion of bone marrow cells caused by a series of somatic mutations (e.g., *DNMT3A, TET2, ASXL1*, and *JAK2*) resulting in pre-malignant clonal expansion of blood cells, is recognized as a major risk factor for atherosclerotic CVD through immune cell modulation of plaque inflammation and stability(*55*). Recent work showed that aged bone marrow-derived cells recruited to atherosclerotic plaques could promote SMC expansion and atherosclerosis(*56*), suggesting that complex cell-cell interactions, including those between different cell types carrying somatic mutations, may regulate plaque vulnerability and clinical CVD events. Therefore, a systematic evaluation of somatic mutations in non-myeloid cells (e.g., SMC/SDCs, ECs, fibroblasts) in atherosclerotic lesions is required to reveal if and to what extent such somatic mutations can increase plaque vulnerability and clinical CVD outcomes and inform the development of new strategies to prevent and cure the disease.

Several lines of evidence, including ours, suggest that certain anti-cancer strategies, such as enhancing macrophage phagocytosis (e.g., anti-CD47 antibody(*57*)) and inhibiting cell proliferation (e.g., ATRA(*6*)), may be atheroprotective. Here we provide proof of principle that niraparib, a chemotherapy targeting DNA damage repair, can both attenuate development of atherosclerosis and induce regression of established disease. Enhancing protective functions of other immune cells, such as macrophage phagocytosis, also seems to attenuate atherosclerosis(*58*). However, some cancer treatment strategies, such as immunotherapy via blocking PD1/PD-L1, can increase risk of CVD events by stimulating cytotoxic T cells to secret atherogenic cytokines(*59*). Conflicting findings for cancer treatments likely reflect the distinct effects of specific cell-targeting (e.g., T-cells, macrophages and SMC/SDCs) on atherosclerosis and risk of clinical complications.

In summary, the complexity of atherosclerosis pathogenesis, as well as SMC tumor-like behaviors and cellular crosstalk among these transformed cell types, requires a comprehensive and thoughtful “athero-oncology” perspective to drive mechanism-based translation and new therapeutic opportunities. A focus on genetic and molecular pathways shared between the two complex diseases while defining the distinctive molecular and cellular process in atherosclerosis can reveal which paradigms of cancer therapy represent new opportunities for targeted interventions for atherosclerotic CVD, particularly in patients with established diseases and already on lipid lowering therapies.

## ACKNOWLEDGMENTS

We thank Theresa C. Swayne for the assistance of microscopy and image analysis and Caisheng Lu and Wan-I Kuo for the aid of flow cytometry analysis and FACS sorting. We thank Dajiang Sun, Tingting Li, and Andy Zhu for the help of H&E staining and slide scanning.

## Author contributions

Conceptualization: HP, MPR

Methodology: HP, CX, MPR

Investigation: HP, SEH, CX, JC, FL, RAS, ESC

Visualization: HP, SEH

Funding acquisition: HP, MPR

Project administration: HP

Supervision: HP, MPR

Writing – original draft: HP, LSR, MPR

Writing – review & editing: all authors

## Data and materials availability

All data and materials used in the study are available in the main text or supplementary materials. Genomic data are publicly available at the GEO (accession number GSE***).

## SUPPLEMENTARY MATERIALS

Materials and Methods

Figs. S1 to S8

