## Supplementary material for "Atherosclerosis is a smooth muscle cell-driven tumor-like disease": Methods and Supplementary Figures

**This PDF file includes:**

Materials and Methods

Figs. S1 to S8

### Materials and Methods

Mice

All mouse experiments were approved by Columbia University Institutional Animal Care and Use Committee (IACUC). *ROSA26*$^{LSL-ZsGreen1}$ (Stock No: 007906), *Ldlr*$^{-/-}$ (Stock No:002207), *Myh11-CreER*$^{T2}$ (Stock No: 019079), and *Kras*$^{LSL-G12D/+}$ (Stock No: 032435) mice were obtained from the Jackson Laboratory and bred onto the C57BL/6J (Stock No: 000664) background. SMC lineage tracing *ROSA26*$^{LSL-ZsGreen1/+}$; *Ldlr*$^{-/-}$; *Myh11-CreER*$^{T2}$ mice were generated as previously described. *Kras*$^{+/+}$; *ROSA26*$^{LSL-ZsGreen1/+}$; *Ldlr*$^{-/-}$; *Myh11-CreER*$^{T2}$ and *Kras*$^{LSL-G12D/+}$ *ROSA26*$^{LSL-ZsGreen1/+}$; *Ldlr*$^{-/-}$; *Myh11-CreER*$^{T2}$ mice were generated by crossing *Kras*$^{LSL-G12D/+}$ mice with SMC lineage tracing mice as indicated in fig. S6A. To induce SMC-specific expression of ZsGreen1, 6-week-old SMC lineage tracing mice were fed tamoxifen diet (Envigo, TD.130859) for one week, then chow diet for another week. Afterwards, mice at the age of 8 weeks were fed western diet (WD, Envigo, TD.88137) for various timepoints as indicated. Of note, as BAC transgene containing *Myh11* promoter-driven *CreER*$^{T2}$ was integrated into the Y chromosome, all *Myh11-CreER*$^{T2}$-relevant mice used in the study are male.

Immunohistochemistry staining

Mouse brachiocephalic artery (BCA) tissues were fixed and frozen blocks were prepared and sectioned as previously described(*6*). Frozen sections (10-μm thickness) were permeabilized with 0.4% Triton X-100/PBS, blocked with 5% BSA, and incubated with primary antibodies at 4°C, overnight, followed by staining of nuclei with Hoechst 33342 (Invitrogen, H3570) and secondary antibodies for 1 hour at RT. After washing with PBS three times, sections were mounted with ProLong Glass Antifade Mountant (Invitrogen, P36984).

Human atherosclerotic arteries were obtained from patients undergoing carotid endarterectomy. Studies with human samples were approved by the Institutional Review Board (IRB) of Columbia University Irving Medical Center. All patients were fully informed of the research and signed IRB-approved informed consent forms. Human atherosclerotic arteries were freshly frozen and sectioned to 10-μm thickness. Human artery sections were fixed with 4% Paraformaldehyde (PFA) for immunohistochemistry staining. The procedure is the same as that with mouse sections.

The following primary antibodies were used for immunostaining: biotin anti-8-OHdG (Abcam, ab183395, 1:200), rabbit anti-4HNE (Alpha Diagnostic, HNE11-S, 1:100), and Phospho-Histone H2A.X (Ser139) (Cell Signaling Technology, 9718, 1:100). The secondary antibodies were as follows: Alexa Fluor 594-conjugated Streptavidin (Molecular Probes, S32356) and Goat anti-Rabbit IgG (H+L) Highly Cross-Adsorbed Secondary Antibody, Alexa Fluor 594 (Invitrogen, A-11037). Immuno-stained sections were visualized using Nikon Eclipse Ti-E inverted microscope at 10× and 20× objective magnifications. The confocal pinhole was set at 1 Airy unit, to produce an optical section of approximately 0.5 μm. Nuclei, ZsGreen1, 8-OHdG/4-HNE/γH2A.X were imaged sequentially with excitation at 405, 488 and 561, respectively, and emission was collected with standard emission filters. Images were analyzed with Fiji/ImageJ software (NIH).

Comet assay

Aortic single cells from SMC lineage tracing mice were prepared as previously reported(*6*). Freshly sorted ZsGreen1$^{+}$ SMCs (from healthy aortas of 8-week-old mice on 0-week WD) and SDCs (from atherosclerotic aorta of mice on 26-week WD) were subject to comet assay using the Comet Assay Single Cell Gel Electrophoresis Assay kit (R&D Systems, 4250050K) according to the manufacturer's manual. The nuclei were stained with SYTOX Green (Molecular Probes). Slides were randomly scanned using Nikon Ti-S microscope and images were analyzed with Fiji/ImageJ software (NIH).

Isolation and *ex vivo* culture of SMCs and SDCs from mouse aortas

SMCs and SDCs were isolated from aortas of SMC lineage tracing mice and *ex vivo* cultured according to published protocols(*60, 61*) with some modifications. Briefly, healthy (for SMC isolation) or atherosclerotic (for SDC isolation, 26-week WD) aortas were removed from euthanized *ROSA26*$^{LSL\text{-}ZsGreen1/+}$; *Ldlr*$^{-/-}$; *Myh11-CreER*$^{T2}$ mice and placed in aortic digestion solution (175 U/mL Collagenase Type II and 1.25 U/mL elastase in DMEM) at 37 °C. After 30-minute digestion, the outer adventitia layer was removed, and the remaining aortas were placed into cell culture medium (DMEM + 10% FBS +1% penicillin/streptomycin) and incubated in 37 °C, 5% $CO_2$ incubator for 2 hours. Then the aortas were cut to into 1-2 mm wide rings and digested in aortic digestion solution for 1 hour. Cells were then transferred to cell culture medium and cultured in 37 °C, 5% $CO_2$ incubator. After cells were expanded for 2 passages, fluorescence-activated cell sorting (FACS) was performed to purify ZsGreen1$^+$ SMCs and SDCs. Primary SMCs within 6 passages were used for experiments, unless indicated.

<u>SA-β-Gal staining</u>

SA-β-Gal staining of culture cells were performed at various passages as indicated using the Senescence β-Galactosidase Staining Kit (Cell Signaling Technology, 9860) based on the manufacturer's protocol. For artery tissue staining, the procedure was slightly modified. Briefly, mouse aortas were fixed in Fixative Solution for 30 min at room temperature followed by incubation in β-Galactosidase Staining Solution at 37°C overnight. Then, BCA tissue were frozen and sectioned into 10-µm sections. Images were captured by Nikon Ti-S microscope and analyzed using Fiji/ImageJ software (NIH).

<u>Cell population doubling level calculation</u>

SMCs and SDCs harvested at each passage (after passage 3) were counted and 50,000 cells/well were then seeded to 6-well plate. Cell population doubling levels were calculated using the published formula(*62*): $PDL_n = PDL_{n-1} + 3.322 (\log C_n - \log C_i)$, in which $PDL_n$ = population doubling level at passage n, $PDL_{n-1}$ = population doubling level at passage n-1, $C_i$ = initial cell number seeded, $C_n$ = yield cell number at passage n.

<u>5-ethynyl-2'-deoxyuridine (EdU) cell proliferation assay</u>

EdU incorporation assay was performed to evaluate cell proliferation rate based on DNA synthesis(*29*). SMCs and SDCs were treated with EdU at 10 µM for 2 hours and 24 hours and DNA synthesis was determined with the Click-iT Plus EdU Cell Proliferation Imaging Kit, Alexa Fluor 647 dye (Invitrogen, C10640) according to the manufacturer's guidance.

<u>Colony formation assay</u>

Cell colony formation assay was performed according to a published protocol(*63*) with some modifications. SMCs and SDCs were seeded to 6-well plate at the density of 200 cells/well. Culture medium was changed every other day. After 10-day culture, cells were stained with 0.5% crystal violet (Sigma-Aldrich, V5265).

<u>Cell invasion assay</u>

Cell invasion assay was conducted based on a previous report(*64*). Transwell inserts (Corning, 3422) were coated with 2.0 mg/mL Matrigel Membrane Matrix (Corning, 356234) in advance. SMCs or SDCs were suspended in DMEM basal medium and seeded at a density of 10,000/insert. Inserts were then placed into cell culture plate with DMEM + 20% FBS. After 24 hours, top of the membrane (including Matrigel and cells) were carefully removed with cotton tipped applicators. Inserts were placed onto slides for imaging.

<u>3-dimensional (3D) tumor spheroid assay</u>

Previous protocols (*65, 66*)were applied and slightly modified for the 3D spheroid formation assay with SMCs and SDCs. SMCs or SDCs were seeded into Costar Ultra-Low Attachment Microplate (Corning, 3473) with a density of 10,000/well and cultured with 3D Tumorsphere Medium XF (Promocell, C-28070) for 10 days. Half of medium was changed every other day.

<u>Cell apoptosis assay</u>

SMCs and SDCs were treated with or without 25 ng/mL TNFα for 24 hours before detecting apoptosis using Annexin V Apoptosis Detection Kit APC (eBioscience, 88-8007-72) through flow cytometry analysis according to the manufacturer's manual. $PI^{-}$/Annexin $V^{-}$ (quadrant 1, Q1) indicates live cells; $PI^{-}$/Annexin $V^{+}$ (Q2) indicates early apoptotic cells; $PI^{+}$/Annexin $V^{+}$ (Q3) indicates late apoptotic cells; and $PI^{+}$/Annexin $V^{-}$ (Q4) indicates necrotic cells(*67*).

Flow cytometry analysis of cancer stem cell markers

Aortic single cells from SMC lineage tracing mice were prepared as previously reported. Cell pellets were resuspended in FACS buffer with rat anti-mouse CD16/CD32 (eBioscience, 16-0161) at 4°C, 30 min, to block unspecific binding of antibodies to Fc receptors. Subsequently, cells were incubated with DAPI, rat anti-mouse CD24-PE/Cy7 (BioLegend, 101821), and rat anti-mouse/human CD44-APC (BioLegend, 103011) for 20 min at 4°C before flow cytometry analysis using BD Influx instrument.

Immunoblotting

SMCs and SDCs were lysed with 2x Laemmli buffer (Bio-Rad) supplemented with 5% β-mercaptoethanol. After incubation in 95°C for 10 min, proteins were subject to immunoblotting using NuPAGE Gels (ThermoFisher) according to the manufacturer's manual. The primary antibodies used for immunoblotting are as follows:

| ***Antibody*** | ***Vendor*** | ***Catalog #*** | ***Dilution*** |
|---|---|---|---|
| **α-Tubulin** | Cell Signaling Technology | 2144 | 1:1,000 |
| **Akt (pan) (11E7)** | Cell Signaling Technology | 4685 | 1:1,000 |
| **Phospho-Akt (Ser473)** | Cell Signaling Technology | 9271 | 1:1,000 |
| **β-Catenin (D10A8)** | Cell Signaling Technology | 8480 | 1:1,000 |
| **HIF-1α (D1S7W)** | Cell Signaling Technology | 36169 | 1:1,000 |
| **Phospho-Histone H2A.X (Ser139)** | Cell Signaling Technology | 9718 | 1:1,000 |
| **MEK1/2 (D1A5)** | Cell Signaling Technology | 8727 | 1:1,000 |
| **Phospho-MEK1/2 (Ser217/221) (41G9)** | Cell Signaling Technology | 9154 | 1:1,000 |
| **NF-κB p65 (C22B4)** | Cell Signaling Technology | 4764 | 1:1,000 |
| **Phospho-NF-κB p65 (Ser536) (93H1)** | Cell Signaling Technology | 3033 | 1:1,000 |
| **p16 INK4A (E5F3Y)** | Cell Signaling Technology | 29271 | 1:1,000 |
| **p21 Waf1/Cip1** | Cell Signaling Technology | 64016 | 1:1,000 |
| **p44/42 MAPK (Erk1/2) (137F5)** | Cell Signaling Technology | 4695 | 1:1,000 |
| **Phospho-p44/42 MAPK (Erk1/2) (Thr202/Tyr204) (D13.14.4E)** | Cell Signaling Technology | 4370 | 1:1,000 |
| **Phospho-p53 (Ser15)** | Cell Signaling Technology | 9284 | 1:1,000 |
| **Smad2/3 (D7G7)** | Cell Signaling Technology | 8685 | 1:1,000 |
| **Phospho-Smad2 (Ser465/467) (138D4)** | Cell Signaling Technology | 3108 | 1:1,000 |
| **Stat3 (124H6)** | Cell Signaling Technology | 9139 | 1:1,000 |
| **Phospho-Stat3 (Tyr705)** | Cell Signaling Technology | 9131 | 1:1,000 |
| **TRADD** | Cell Signaling Technology | 3694 | 1:1,000 |

| | | | |
|---|---|---|---|
| **TRAF2 (C192)** | Cell Signaling Technology | 4724 | 1:1,000 |
| **Phospho-TRAF2 (Ser11) (E2B6L)** | Cell Signaling Technology | 13908 | 1:1,000 |

The following secondary antibodies were used for immunoblotting analysis: Peroxidase AffiniPure Donkey Anti-Mouse IgG (H+L) (Jackson Immuno Research Labs, 715035151) and Peroxidase AffiniPure Donkey Anti-Rabbit IgG (H+L) (Jackson Immuno Research Labs, 711035152).

CNV analysis

To identify possible copy number variations (CNVs) during atherosclerosis progression, we applied inferCNV(*24*), a program that estimates large scale genomic copy number profiles, and CopyKAT(*23*), a copy number karyotyping method using Bayesian segmentation approach on mouse and human scRNA-seq. The following parameters were used in inferCNV analysis: cutoff=0.1, cluster_by_groups=TRUE, denoise=TRUE, HMM=TRUE. The following parameters were used in CopyKAT analysis: id.type="S", cell.line="no", LOW.DR=0.05, UP.DR=0.2. A reference set was defined in both inferCNV and CopyKAT analyses. For all mouse CNV analyses, ZsGreen1$^+$ SMCs from mice on 0 week of western diet were used as reference set(*6*). In mouse atherosclerosis progression dataset, ZsGreen1$^+$ SMCs, minor SMCs, SEMs, fibrochondrocytes, fibroblasts, and macrophages from mice on 16 weeks of western diet were compared to the reference set. For human CNV analysis, T cells from each scRNA-seq database were used as reference set to minimize the number of CNVs. SMCs, intermediate cell state, fibrochondrocytes, fibroblasts, and macrophages from each subject were compared to the reference.

Relative expression from inferCNV output was log-transformed and visualized on heatmaps. The log of relative expression from CopyKAT was plotted on heatmaps directly. As the CNV signal is small and the log ratios center around zero across loci, we performed Levene's test on log ratio of expression to assess the variance between populations. Briefly, the median log ratio is calculated at each locus across cells in a population to get locus level CNV. The absolute difference between locus level CNV and their mean is then compared between populations. Tukey's test was used for multiple comparison. Two populations were considered to have significant difference in the variance of CNV signal if the adjusted *P*-value is <0.05.

PROGENy analysis

PROGENy builds models of pathway responsive gene expression from publicly available perturbation experiments and infers pathway activity using a linear model(*33*). To investigate cancer-related pathway activity alteration during atherosclerosis progression, we applied PROGENy on scRNA-seq of mouse on 16 weeks of western diet using built-in models of 12 pathways. Specifically, normalized counts from ZsGreen1+ SMCs, SEM cells, and fibrochondrocytes were used to infer pathway activity using the top 500 genes from each pathway model. Kruskal-Wallis test was performed to compare ranks of pathway scores across populations. Dunn's test was used for post-hoc comparison where scores from SEMs and fibrochondrocytes were compared to SMCs. *P*-values were adjusted by Benjamini-Hochberg approach.

Serum cholesterol measurement

Mouse blood was collected via retro-orbital sinus plexus into BD Vacutainer Venous Blood Collection Tubes (BD 367812) and left at room temperature for 2-3 hours followed by centrifuging at 1,000x g for 15 min to obtain serum. Total serum cholesterol concentration was measured using the Total Cholesterol E Kit (FUJIFILM Medical Systems, 99902601) based on the manufacturer's instruction.

*In vivo* studies of niraparib effects on atherosclerosis

*Administration of niraparib during progression of atherosclerosis:* After induction with tamoxifen and feeding WD for 4 weeks, *ROSA26*$^{LSL\text{-}ZsGreen1/+}$; *Ldlr*$^{-/-}$; *Myh11-CreER*$^{T2}$ mice were orally administrated with vehicle (corn oil, Sigma-Aldrich, C8267) or niraparib (Selleck Chemicals, S2741, 10 mg/kg mice) 3 times per week for 12 weeks. During this time, the mice were continued on WD and sacrificed after 16-week WD feeding for analysis.

*Niraparib treatment in mice with established atherosclerosis:* 16-week WD fed *ROSA26*$^{LSL-ZsGreen1/+}$; *Ldlr*$^{-/-}$; *Myh11-CreER*$^{T2}$ mice were continued on WD and orally fed with vehicle or niraparib 3 times per week as described above for additional 8 weeks. After that, mice were sacrificed for further analysis.

Hematoxylin and eosin (H&E) staining

Apical parts of mouse hearts were collected and fixed in Histochoice Tissue Fixative for 3 hours before embedding in Tissue Frozen Medium. H&E staining was performed using aortic sinus sections and slides were scanned in the Histology Service of the Molecular Pathology Shared Resource at Columbia University Irving Medical Center. Atherosclerotic lesion area, necrotic area, and fibrous cap were measured using Aperio ImageScope software (Leica).

*En face* Oil Red O staining

Preparation of *en face* aorta and staining were conducted as previously described(*68*). Whole aortas were isolated from sacrificed mice and all surrounding adipose tissues were removed. Clean aortas were fixed in 4% PFA at 4°C overnight, cut longitudinally, and pinned on dissecting dishes with lumen side up. *En face* aortas were then stained with 0.25% ORO for 15min, visualized under the stereomicroscope (Leica, M80), and captured using camera (Leica).

Quantification and statistical analysis

Statistical analysis was performed using GraphPad Prism V9 software (GraphPad).

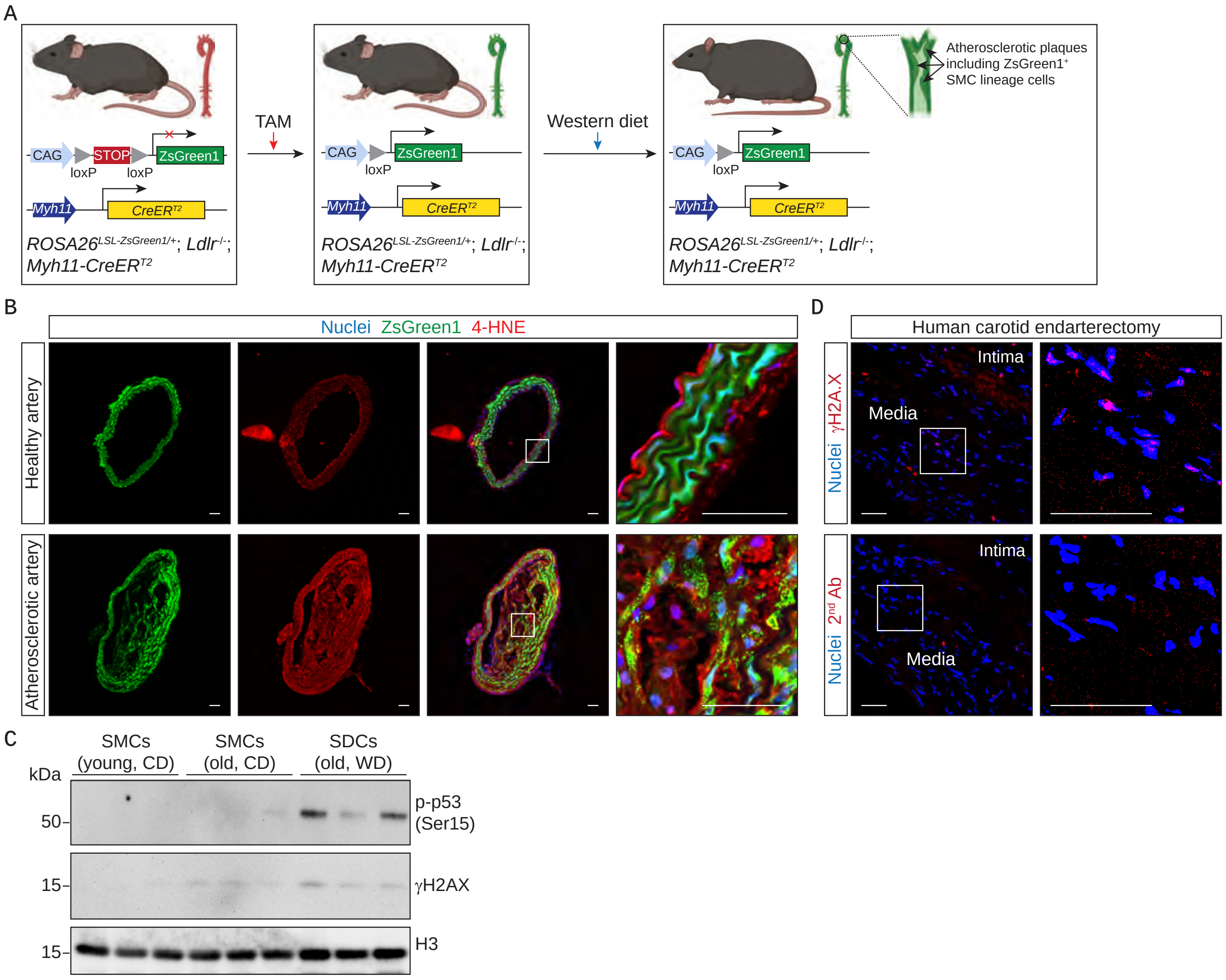


**Fig. S1. Oxidative stress and enrichment of DNA damage-related proteins in mouse SMC lineage cells and human atherosclerosis.** (**A**) Schematic showing induction of ZsGreen1 expression in vascular SMCs in SMC linage tracing *ROSA26*$^{ZsGreen1/+}$; *Ldlr*$^{-/-}$; *Myh11-CreER*$^{T2}$ mice with tamoxifen (TAM) and induction of atherosclerosis with western diet (WD). (**B**) Representative images of lipid peroxidation marker, 4-Hydroxy-2-Nonenal (4-HNE), staining in brachiocephalic artery (BCA) sections from mice on 0-week (healthy artery) and 16-week (atherosclerotic artery) WD. (**C**) Western blot of phospho-p53 (Ser15) and γH2A.X with protein extracts from primary SMCs and SDCs. SMCs were isolated from aortas of young (8-week-old) and old (34-week-old) mice on chow diet (CD). SDCs from atherosclerotic aortas of mice on WD for 26 weeks (old, WD). (**D**) Immunohistochemistry (IHC) staining of γH2A.X with human carotid atherosclerotic sections. Secondary antibody only (2$^{nd}$ Ab) was used as a negative control for staining. Scale bars, 50 μm.

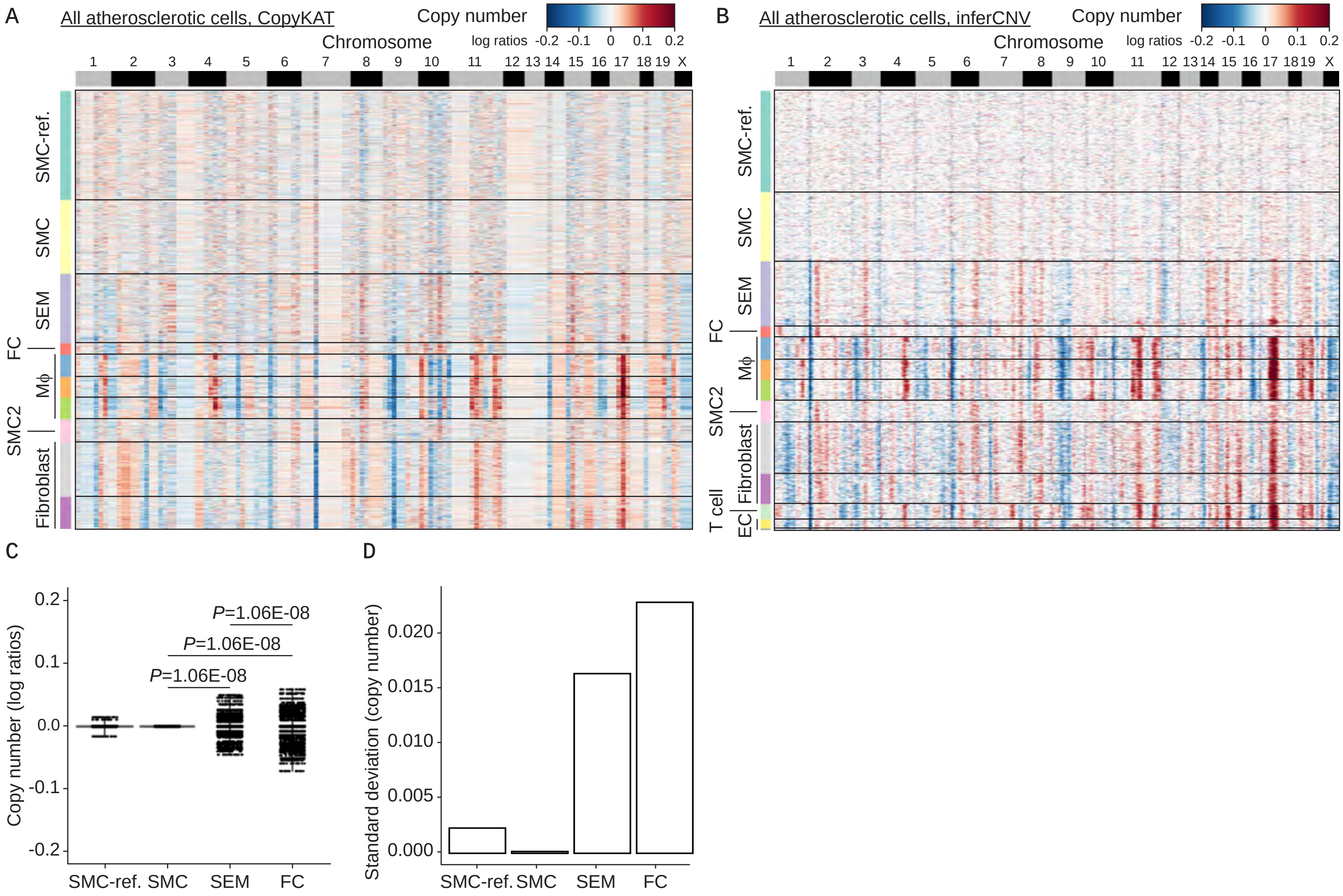


**Fig. S2. CNV analysis of mouse atherosclerosis scRNA-seq database.** (**A**) Clustered heat map showing copy number profiles estimated by CopyKAT in all atherosclerotic cells from scRNA-seq database of 16-week WD fed mice. SMCs from 0-week WD fed mice were as reference (SMC-ref.). Macrophage (Mϕ). (**B**) Clustered heat map showing copy number profiles estimated by inferCNV in all atherosclerotic cells from the same scRNA-seq database in (**A**). (**C**) Analysis of copy numbers (shown as log ratios) in SMC-ref., SMCs, SEM cells, and FCs show increase of CNVs in SEM cells and FCs versus SMC-ref. and SMCs. (**D**) Standard deviation of log ratios (copy number) is increased in SEM cells and FCs versus SMC-ref. and SMCs.

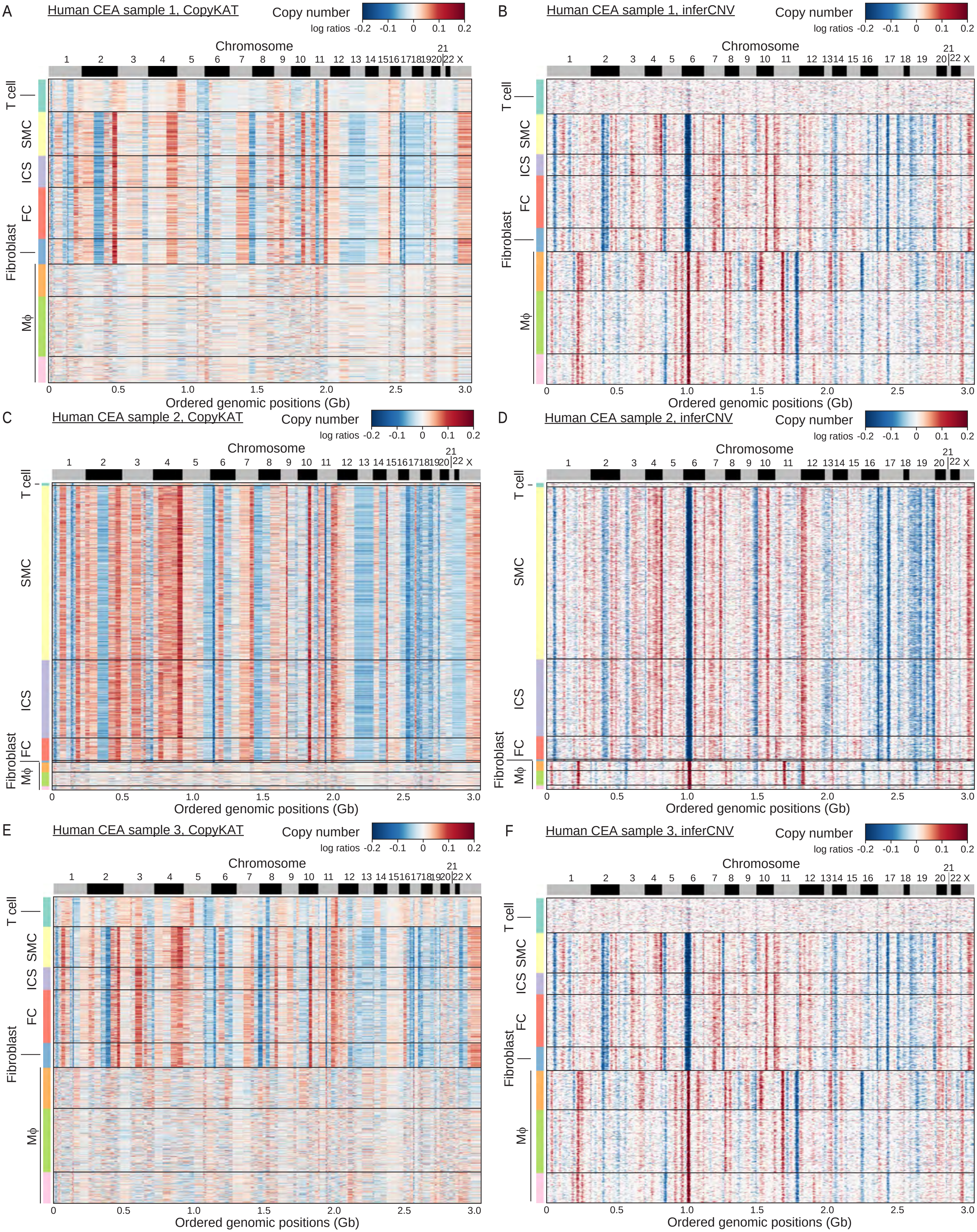
A
Human CEA sample 1, CopyKAT
B
Human CEA sample 1, inferCNV
C
Human CEA sample 2, CopyKAT
D
Human CEA sample 2, inferCNV
E
Human CEA sample 3, CopyKAT
F
Human CEA sample 3, inferCNV
Copy number
log ratios -0.2 -0.1 0 0.1 0.2
Chromosome
1 2 3 4 5 6 7 8 9 10 11 12 13 14 15 16 17 18 19 20 21 22 X
T cell
SMC
ICS
FC
Fibroblast
Mϕ
0 0.5 1.0 1.5 2.0 2.5 3.0
Ordered genomic positions (Gb)

**Fig. S3. CNV analysis of multiple human atherosclerosis scRNA-seq databases.** (**A**-**F**) Clustered heat map showing copy number profiles estimated by CopyKAT (**A**, **C**, and **E**) or inferCNV (**B**, **D**, and **F**) in human atherosclerotic cells from scRNA-seq databases of human CEA sample 1 (**A** and **B**), 2 (**C** and **D**), and 3 (**E** and **F**). Human atherosclerotic plaque cells include T cells, SMC, intermediate cell state (ICS), fibrochondrocyte (FC), fibroblast, and macrophage (M$\phi$).

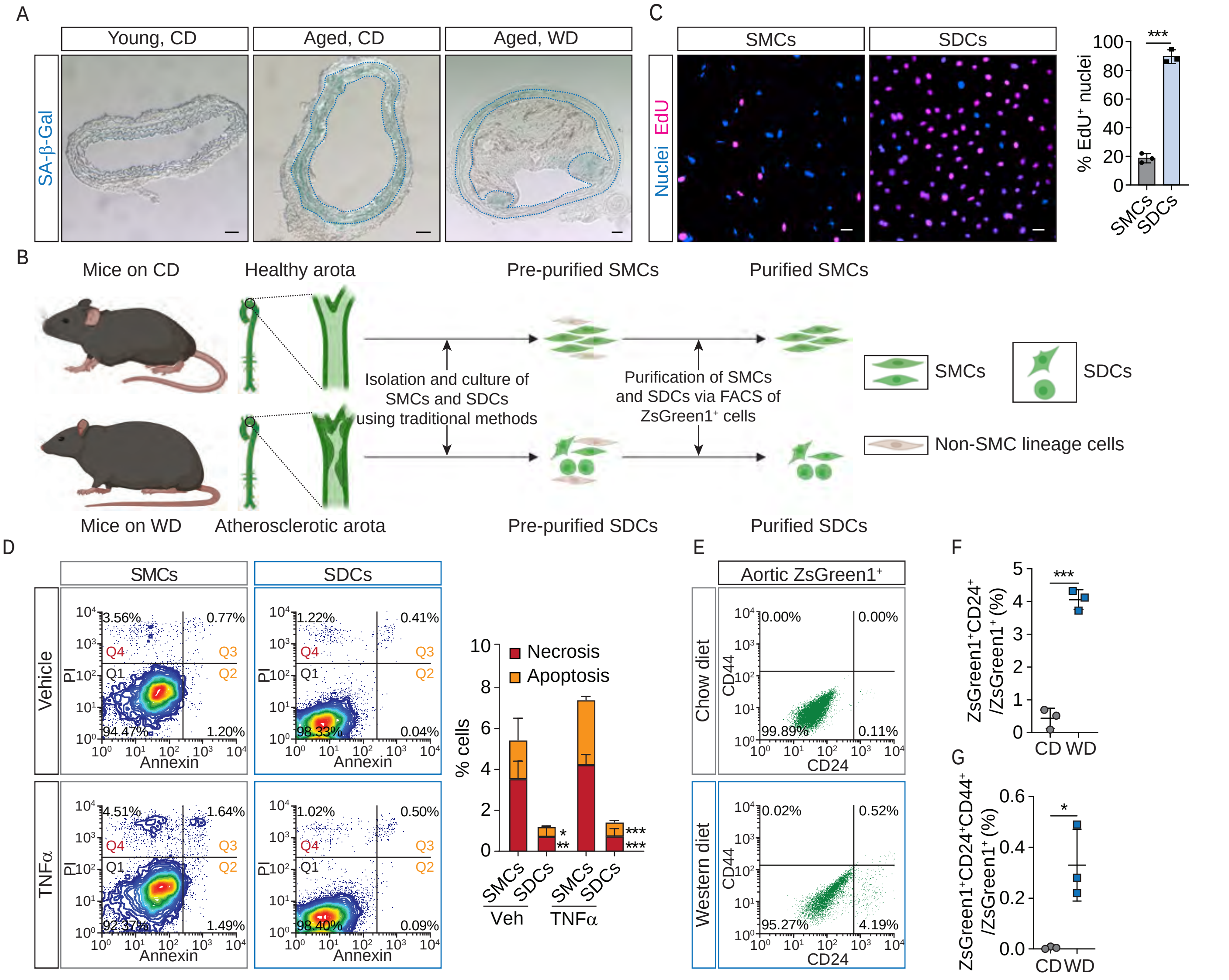


**Fig. S4. SMC-derived cells harbor multiple tumor cell-like characteristics.** (**A**) Staining of senescence-associated beta-galactosidase (SA-β-Gal), a biomarker of cellular senescence, in mouse BCA sections from young (12-week-old) and aged (34-week-old) mice (on regular chow diet (CD), without atherosclerosis) and atherosclerotic mice (aged, on 22-week WD). Scale bars, 50 µm. (**B**) Workflow of isolation, culture, and purification of SMCs and SDCs from mouse aortas. (**C**) SMCs and SDCs were incubated with 5-ethynyl-2'-deoxyuridine (EdU) for 24 hours. Click-iT EdU assay indicated that SDCs had much higher proliferative rate (proportion of $EdU^+$ nuclei) than SMCs. (**D**) Flow cytometry analysis of apoptosis and necrosis of SMCs and SDCs treated with vehicle (Veh) or TNFα. $PI^-$/Annexin $V^-$ (quadrant 1, Q1) indicates live cells; $PI^-$/Annexin $V^+$ (Q2) indicates early apoptotic cells; $PI^+$/Annexin $V^+$ (Q3) indicates late apoptotic cells; and $PI^+$/Annexin $V^-$ (Q4) indicates necrotic cells. (**E**) Flow cytometry analysis of cancer stem cell surface markers, CD24 and CD44, in $ZsGreen1^+$ SMCs and SDCs freshly isolated from CD-fed mice (8-week-old) and WD-fed mice (26-week WD), respectively. (**F** and **G**) Statistical analysis of proportion of $ZsGreen1^+CD24^+/ZsGreen1^+$ and $ZsGreen1^+CD24^+CD44^+/ZsGreen1^+$. Significance was determined by unpaired two-tailed t test. N=3, *$P$<0.05, **$P$<0.01, ***$P$<0.001.

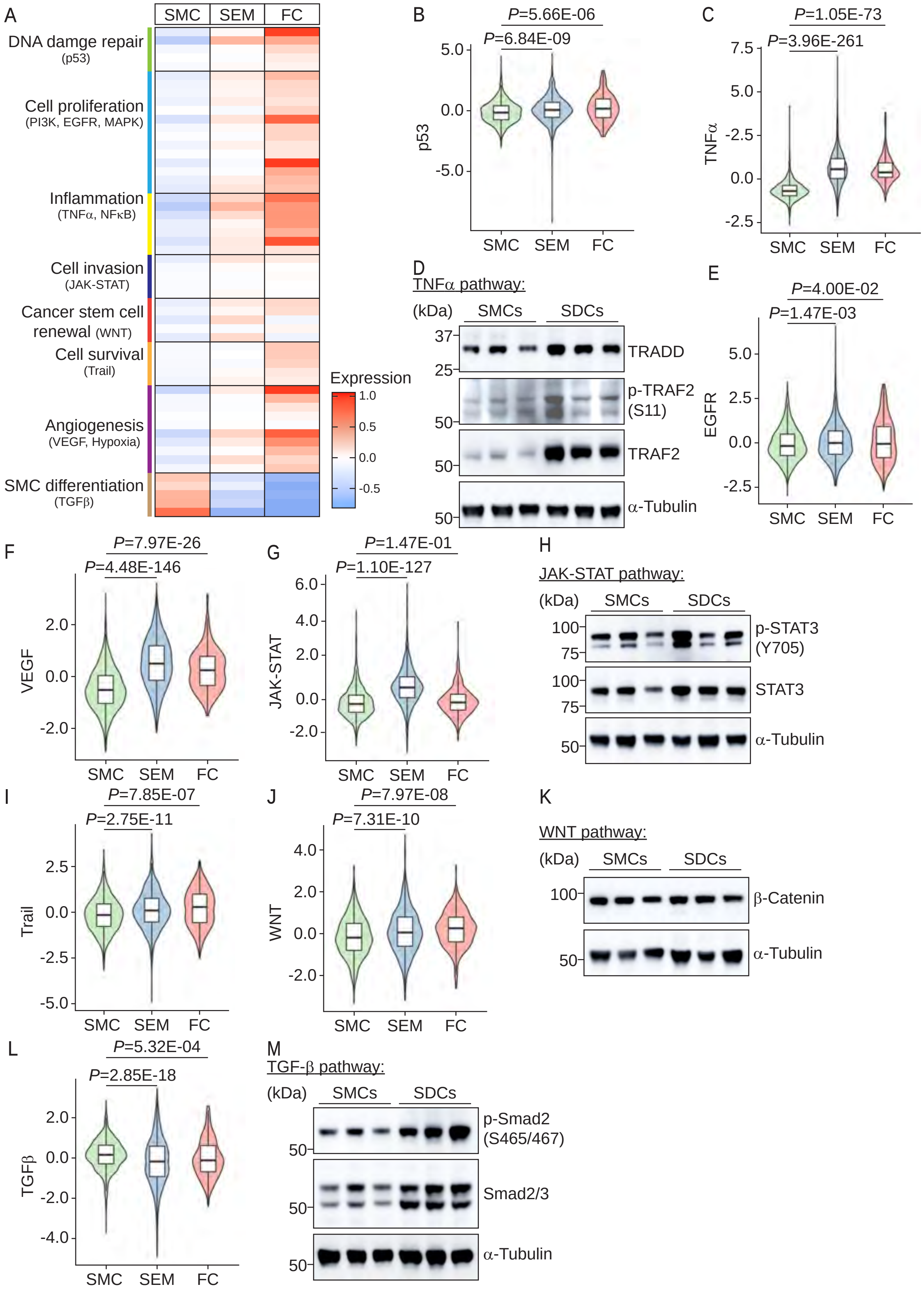
A
SMC
SEM
FC
DNA damge repair
(p53)
Cell proliferation
(PI3K, EGFR, MAPK)
Inflammation
(TNFα, NFκB)
Cell invasion
(JAK-STAT)
Cancer stem cell renewal (WNT)
Cell survival
(Trail)
Angiogenesis
(VEGF, Hypoxia)
SMC differentiation
(TGFβ)
Expression
1.0
0.5
0.0
-0.5
B
P=5.66E-06
P=6.84E-09
p53
C
P=1.05E-73
P=3.96E-261
TNFα
D
TNFα pathway:
(kDa)
SMCs
SDCs
TRADD
p-TRAF2 (S11)
TRAF2
α-Tubulin
E
P=4.00E-02
P=1.47E-03
EGFR
F
P=7.97E-26
P=4.48E-146
VEGF
G
P=1.47E-01
P=1.10E-127
JAK-STAT
H
JAK-STAT pathway:
p-STAT3 (Y705)
STAT3
I
P=7.85E-07
P=2.75E-11
Trail
J
P=7.97E-08
P=7.31E-10
WNT
K
WNT pathway:
β-Catenin
L
P=5.32E-04
P=2.85E-18
TGFβ
M
TGF-β pathway:
p-Smad2 (S465/467)
Smad2/3

**Fig. S5. Activated cancer-associated signaling pathways in SDCs versus SMCs.** (**A**) Heat map showing top 10 genes related to each of 12 cancer-associated signaling pathways in SMCs, SEM cells, and FCs estimated by Pathway RespOnsive GENes (PROGENy). scRNA-seq data of ZsGreen1$^+$ cells from 16-week WD fed mice were used for analysis. (**B** and **C**) Violin plots show PROGENy scores for p53 (**B**) and TNFα (**C**) pathways. (**D**) Immunoblotting results indicate that key transducers of TNFα signaling, TRADD and phospho-TRAF2 (S11), were increased in SDCs versus SMCs. (**E**-**G**) Violin plots show PROGENy scores for EGFR (**E**), VEGF (**F**), and JAK-STAT (**G**) pathways. (**H**) Immunoblotting results indicate that a key transducer of JAK-STAT signaling, phospho-STAT3 (Y705) was increased in SDCs versus SMCs. (**I** and **J**) Violin plots show PROGENy scores for Trail (**I**) and WNT (**J**) pathways. (**K**) Immunoblotting showing the protein level of β-Catenin, a key transducer of WNT signaling in SMCs and SDCs. (**L**) Violin plot shows PROGENy scores for TGFβ pathway. (**M**) Immunoblotting results indicate that phospho-Smad2 (S465/467), a key transducer of TGFβ signaling, was increased in SDCs versus SMCs. *P* values are indicated.

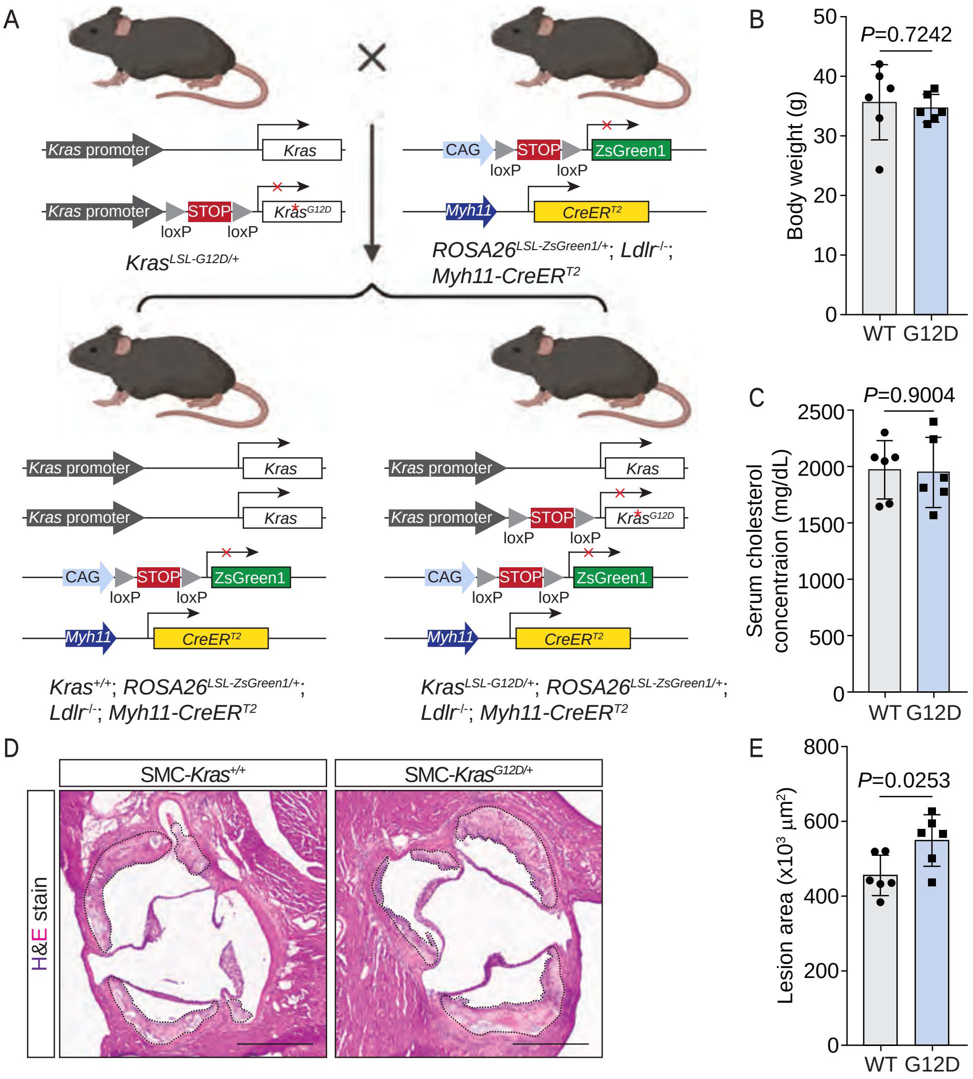


**Fig. S6. SMC-*Kras*$^{G12D}$ exacerbates atherosclerosis.** (**A**) Schematic of generation of $Kras^{+/+}$; $ROSA26^{LSL-ZsGreen1/+}$; $Ldlr^{-/-}$; $Myh11$-$CreER^{T2}$ (SMC-$Kras^{+/+}$) and $Kras^{LSL-G12D/+}$; $ROSA26^{LSL-ZsGreen1/+}$; $Ldlr^{-/-}$; $Myh11$-$CreER^{T2}$ (SMC-$Kras^{G12D/+}$) mice. (**B** and **C**) Body weight (**B**) and serum cholesterol levels (**C**) of SMC-$Kras^{+/+}$ (wild type, WT) and SMC-$Kras^{G12D/+}$ (G12D) mice at 16-week WD. N=6 mice/group. (**D**) Representative images of hematoxylin and eosin (H&E)-stained aortic sinus sections from SMC-$Kras^{+/+}$ (WT) and SMC-$Kras^{G12D/+}$ (G12D) mice after 16-week WD. Lesion areas were indicated with black dotted lines. (**E**) Statistical analysis of lesion areas in aortic sinus sections. Scale bars, 500 µm. Significance was determined by unpaired two-tailed t test. N=6 mice/group. *P* values are indicated.

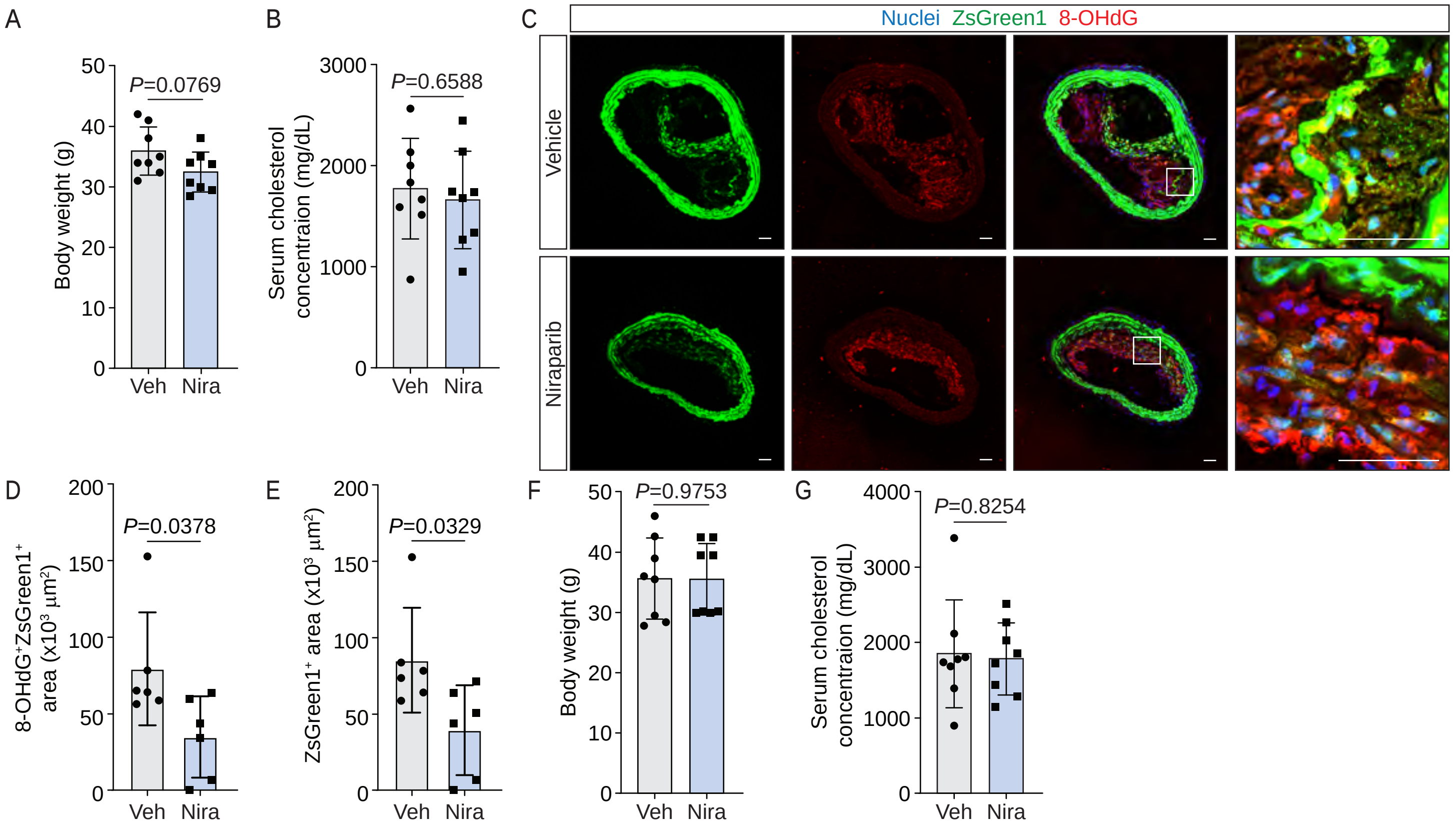


**Fig. S7. Effects of niraparib on mice in atherogenesis.** (**A** and **B**) Body weight (**A**) and serum cholesterol concentration (**B**) of vehicle (Veh) and niraparib (Nira)-treated mice at 16-week WD. N=8 mice/group. (**C**) Representative images of 8-OHdG-stained BCA sections from vehicle and niraparib-treated mice at 16-week WD. (**D** and **E**) Statistical analysis of 8-OHdG$^{+}$ZsGreen1$^{+}$ areas (**D**) and ZsGreen1$^{+}$ areas in neointima (**E**) of BCA sections from Veh and Nira-treated mice at 16-week WD. N=6 mice/group. (**F** and **G**) Body weight (**F**) and serum cholesterol concentration (**G**) of Veh and Nira-treated mice at 24-week WD. N=8 mice/group. Scale bars, 50 µm. Significance was determined by unpaired two-tailed t test. *P* values are shown.

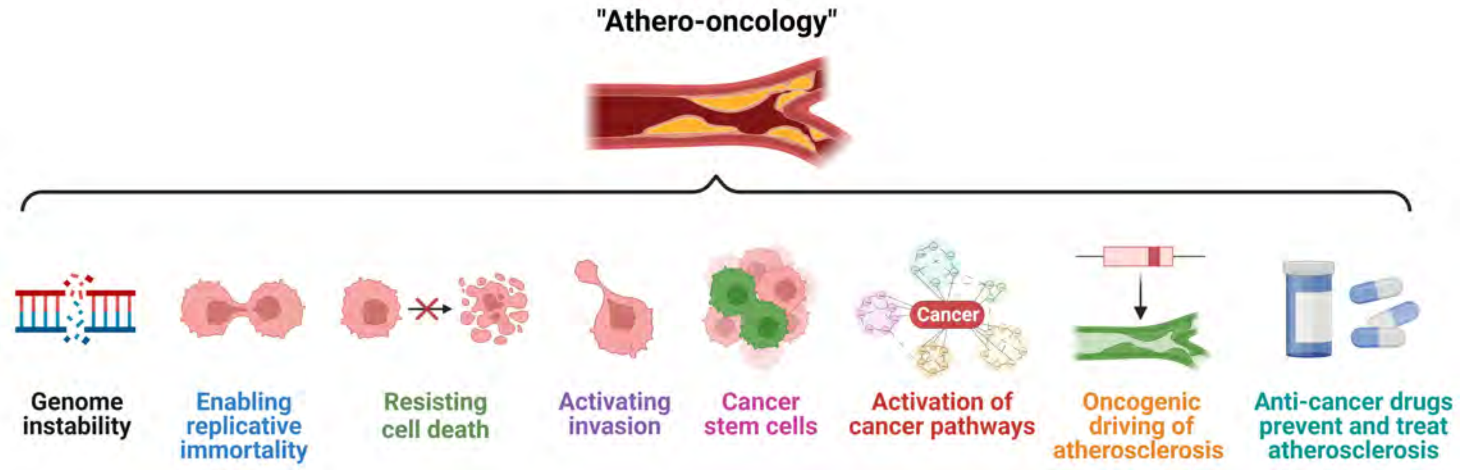


**Fig. S8. A novel concept, "athero-oncology", indicates that atherosclerosis is a tumor-like disease.** SDCs in atherosclerosis show multiple characteristics of tumor cells, including genomic instability, enabling replicative immortality, resisting cell death, invasiveness, and cancer stem cell-like features. Extensive cancer-associated gene regulatory networks are activated in SDCs. An oncogenic mutation, *Kras*$^{G12D}$, accelerates SMC phenotypic switching and exacerbates atherosclerosis. Chemotherapies for cancer treatment (e.g., niraparib) may also be beneficial for prevention and treatment of atherosclerotic cardiovascular disease.
